# Molecular and cell type-specific determinants of inferior colliculus development and auditory function

**DOI:** 10.64898/2026.07.18.739349

**Authors:** Miranda Jankovic, Yuan-Tai Huang, Andrew Holley, Sophia Kirkland, Ashwinikumar Kulkarni, Hitomi Sakano, Jay R. Gibson, Genevieve Konopka

**Affiliations:** Department of Neuroscience, UT Southwestern Medical Center, Dallas, TX 75390, USA; Department of Neurobiology, University of California, Los Angeles, Los Angeles, CA 90095, USA

## Abstract

Sound perception requires sensory information processing through brain regions where alterations to cellular composition or function may impact auditory-relevant behaviors. The inferior colliculus (IC) is a central midbrain hub for integrating and transforming auditory information prior to relay to the forebrain. However, the molecular logic that underlies the cell type specification of the IC remains unknown. Here, using a multiomic approach, we define the transcriptional and chromatin landscapes that underlie IC cell type diversity. We identify distinct glutamatergic neuronal subclasses and the gene regulatory programs that drive their specification, maturation, and survival. We show that perturbation to the transcription factor FOXP2, previously implicated in speech and language as well as brain disorders with altered sensory processing, selectively disrupts the specification and survival of three newly defined glutamatergic neuronal subclasses. We link these molecular and cellular disruptions to functional deficits in auditory processing including altered brainstem and forebrain responses and impaired behavioral sensitivity to acoustic stimuli. Together, these results link gene regulatory mechanisms to cell type-specification in the IC, providing insight into how molecular control of neuronal identity in midbrain sensory centers contributes to systems-level processing of sound.

## Introduction

The inferior colliculus (IC) is a midbrain nucleus in the ascending auditory pathway that serves as the major subcortical synaptic relay and a central hub for the integration of auditory and non-auditory information before its transmission to higher sensory centers. Beyond simple signal relay, the IC participates in multiple levels of auditory computation, including temporal and frequency processing, feature extraction, multisensory integration, and bidirectional modulatory control over other auditory nuclei ^1,2^. Through these operations, the IC plays a critical role in shaping how sound information is transformed, represented, and conveyed throughout the auditory system, positioning it as a key determinant of mature auditory function.

The IC comprises three major subdivisions, the central nucleus (CIC), dorsal cortex (DCIC), and lateral cortex (LCIC), each with distinct circuitry and functional roles. The CIC participates in the canonical lemniscal pathway and is organized with fibrodendridic laminae and tonotopically with upstream inputs for precise auditory signal transmission ^3^. In contrast, the LCIC integrates auditory and non-auditory inputs and is associated with non-lemniscal processing ^4^. The DCIC receives strong corticofugal input and is implicated in higher-order auditory modulation ^1^. Tracing studies further demonstrate that molecularly defined IC neuron populations exhibit distinct projection patterns in each region ^5^. The establishment of IC cellular composition and circuitry occurs largely before hearing onset and is subsequently refined during postnatal auditory critical periods ^6^.

Historically, IC neurons have been classified based on their anatomical location, morphology, electrophysiological properties, and a limited number of molecular markers ^4^. However, few of these features can reliably predict correlation with each other ^7^ and as a result, neuronal subtypes within the IC remain poorly resolved ^8,9^. Thus, while prior anatomical and physiological studies have defined IC connectivity and response properties, how the molecular identities and transcriptional programs that specify IC neuronal subtypes and how they contribute to auditory processing and sensory sensitivity are unknown. Improved molecular resolution of IC cell types would provide a much-needed toolkit for functionally dissecting IC circuits, especially during developmentally sensitive critical periods.

Given the central role of the IC as the principal auditory relay from brainstem to forebrain and as a major center for auditory signal processing and multisensory integration ^1^, we hypothesized that transcription factors with cell type-specific expression in the IC with known functional links to auditory function might influence cell type contributions to early auditory computations that shape downstream circuit function and behavior. By implementing a cell type-specific molecular profiling approach, we found robust and specific cell type-specific expression of FOXP2 in a subset of glutamatergic neurons in the IC. FOXP2 is a transcription factor expressed in specific neuronal subtypes throughout the developing and mature brain and regulates gene expression programs essential for neuronal function ^10,11^. Variants in *FOXP2* have been associated with a broad range of neurodevelopmental disorders (NDDs) including attention-deficit/hyperactivity disorder ^12^, schizophrenia ^13^, and autism spectrum disorder (ASD) ^14^, all of which can feature prominent sensory processing comorbidities. *FOXP2* variants are also implicated in childhood apraxia of speech and orofacial dyspraxia, which is frequently accompanied by impairments in auditory-linguistic processing and in the use of auditory feedback to guide vocal output ^15^. These features further highlight the importance of sensory–motor integration in FOXP2-associated pathology ^16–18^.

Although the role of FOXP2 has been extensively studied in cortical ^19–23^, striatal ^22–30^, and cerebellar regions ^22,23,31^, its functional significance in the IC remains unexplored. To examine the contribution of FOXP2 to cell type specific development and overall auditory function, we generated an IC glutamatergic neuron–specific *Foxp2* conditional knockout mouse model and combined behavioral and electrophysiological assays of IC-dependent auditory processing with paired single-nucleus transcriptomic and chromatin accessibility profiling. We found that FOXP2 selectively regulates the identity, survival, and developmental gene programs of defined IC excitatory neuron subclasses. Disruption of these programs altered auditory circuit output and downstream cortical responses, establishing the IC as a critical midbrain locus through which FOXP2 shapes auditory system development and function in ways relevant to NDDs. FOXP2 expression patterns are highly conserved across rodents, non-human primates, and humans, even within homologous brain regions in vocal-learning birds indicating an evolutionarily conserved FOXP2 function across multiple neural systems ^32,33^. Thus, studying FOXP2 function in the relatively conserved auditory system of the mouse may provide insights that broadly applicable across species ^34^.

## Results

### Transcriptomic identification of mouse IC neuronal subtypes by snRNA-seq

To transcriptomically define IC neuronal subclasses and identify the cell subtypes, we generated paired snRNA and snATAC-seq datasets from adult mouse IC. We identified major IC cell types including excitatory neurons, inhibitory neurons, astrocytes, and oligodendrocytes in the IC. Within the neuronal population (*Rbfox3* and *Snap25* positive), glutamatergic neurons are identified by expression of *Slc17a6* (*Vglut2*), while inhibitory neurons are marked by the expression of *Slc32a1* (*Vgat*), *Gad1*, and *Gad2*. Receiver operating characteristic (ROC) analysis was performed to identify the genes whose expression most effectively distinguished individual subclusters. We selected genes with the highest discriminative power, with particular emphasis on transcription factors and neurotransmitter-related genes previously examined in IC studies and identified 7 excitatory and 2 inhibitory clusters in addition to astrocytes and oligodendrocytes (**Fig 1A**).

**Figure 1:**
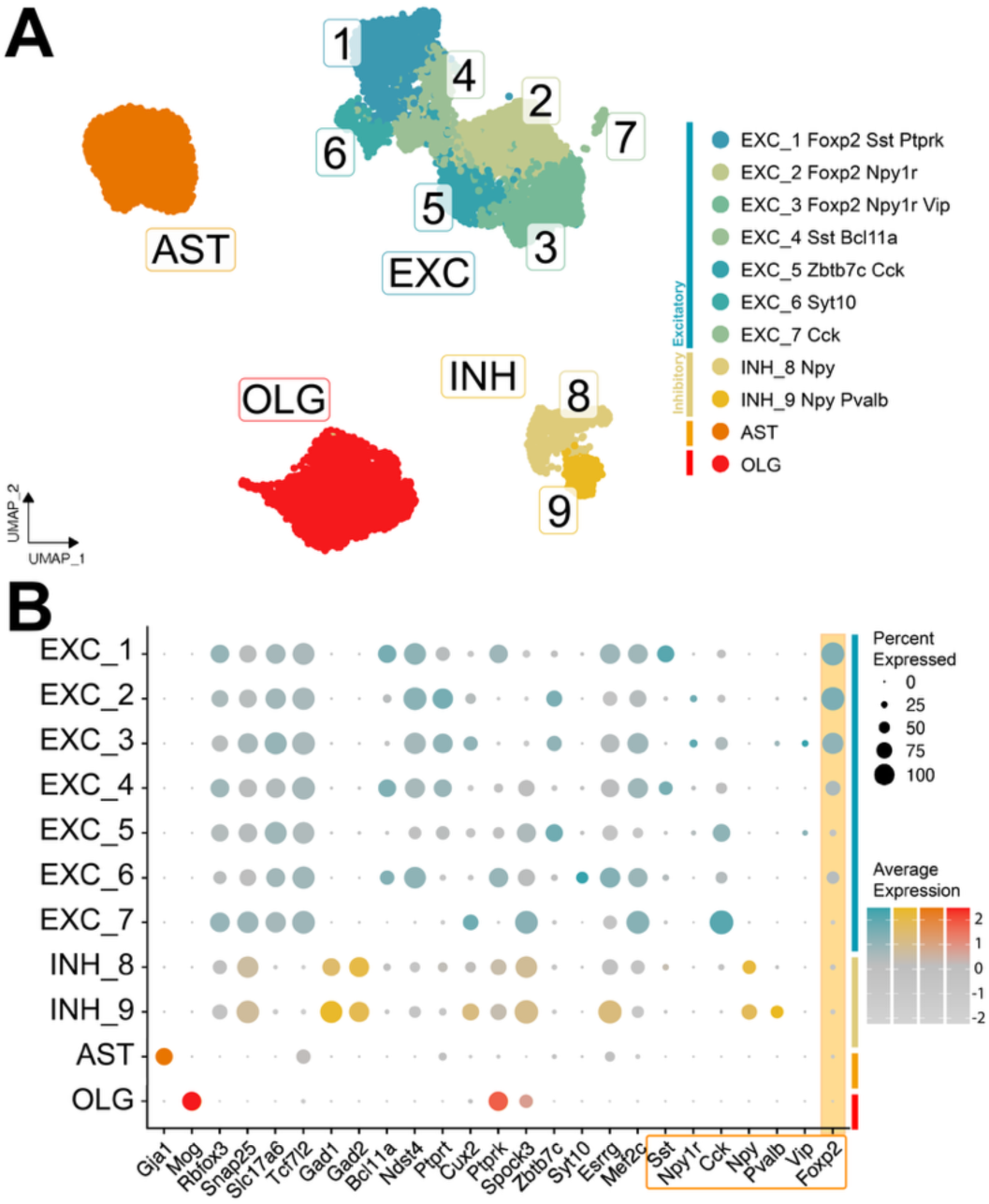
The cellular composition of the IC shows clusters of annotated glutamatergic and GABAergic neurons. (A) UMAP of control data labeled with subtype annotation, colored by major cluster cell type. Annotation numbers correspond with labeled excitatory and inhibitory subtypes. Legend lists extended neuronal subtype cluster names. N = 3 CTL / 3 cKO samples, each sample is pooled from 2 individuals of the same sex, and age range. (B) Dot-plot of the percent nuclei expressing and average expression of genes for each subcluster. Boxed genes denote primary markers of known IC cell types. *Foxp2* expression highlighted in orange.

We used several established neuropeptide markers to delineate distinct neuronal populations (**Fig. 1B**). *Sst* robustly defines specific clusters within excitatory nuclei, EXC 1 and 4. *Cck*, previously used to define a unique IC cell type^37^, shows a more heterogeneous expression pattern, variably marking multiple excitatory subclusters. EXC 7 in particular is strongly positive for *Cck* in contrast. *Vip* expression is detected in a small proportion of cells across select excitatory clusters but does not define a homogeneous subtype^38^. EXC 3 has the largest proportion of *Vip* expressing nuclei, but it still only accounts for a subset of nuclei within the cluster. Among inhibitory neurons, *Pvalb* and *Npy* expression is observed. Cluster INH 8 expresses only *Pvalb,* while INH 9 has overlapping *Pvalb* and *Npy* expression^39^.

Consistent with previous reports, we also observe strong expression of transcription factors known be expressed in the IC such as *Tcf7l2* and *Bcl11a* ^40^, which further delineates glutamatergic subtypes (**Fig. 1B**). Moreover, several neuronal clusters exhibit strong *Foxp2* expression, including excitatory clusters EXC 1, 2, and 3 (**Fig. 1B**). Among these, EXC 1 represents the largest and most homogeneous glutamatergic cluster, identified by a high proportion of neurons in the cluster co-expressing *Foxp2* with robust *Sst* expression. A second *Foxp2*+ excitatory supergroup comprising of EXC 2 and 3 display greater heterogeneity and includes subpopulations expressing *Vip*, *Cck*, and neuropeptide Y receptor type 1 (*Npy1r*). These *Foxp2*+ excitatory clusters are further distinguished by strong correlation with *Ndst4* expression, an enzyme predicted to be involved in proteoglycan biosynthetic processes, and absence of *Spock3*, a calcium-binding proteoglycan protein. Based on previous literature linking FOXP2 to auditory function, we next wanted to assess the role of FOXP2 to IC cellular specification and function.

### IC size and glutamatergic neuron density are regulated by FOXP2

To determine cell type-specific contributions to IC development and function, we generated a conditional knockout of *Foxp2* (*Foxp2* cKO or cKO) in glutamatergic IC neurons by crossing *Foxp2*^fl/fl^ mice with Ntsr1-Cre (GN209) mouse line ^41^. Unlike other Ntsr1-Cre lines, GN209 targets relatively few brain regions and sparse neuronal populations in the brain, but it is selectively expressed across primary sensory auditory structures spanning from the cochlear nuclei to the primary auditory cortices. It strongly targets glutamatergic neurons in the IC ^42^ and is not expressed in other FOXP2 expressing areas such as the neocortex ^43^. Thus, among the primary auditory pathways, Ntsr1-Cre and FOXP2 are most strongly co-expressed in glutamatergic IC neurons, resulting in selective FOXP2 loss-of-function in the IC (**Fig. S1**).

To confirm efficient *Foxp2* deletion in glutamatergic IC neurons, we crossed these mice to Cre reporter lines and performed co-labeled immunohistochemical (IHC) analysis. Samples consisted of mixed male and female adult (postnatal day 50-P60) IC sections, allowing identification of Cre-positive (YFP⁺ or tdTom^+^, both pseudocolored green) glutamatergic neurons across all IC subnuclei. *Foxp2* cKO mice developed without overt gross abnormalities or increased morbidity relative to control littermates, with comparable adult weights (**Fig. S1L**). In control mice, FOXP2⁺ cells were most abundant in the DCIC and CIC nuclei of the IC, with more modest population in the LCIC, mirrored by YFP^+^ cell density (**Fig. 2A,E**). In control ICs, 60% of FOXP2^+^ cells co-expressed YFP (FOXP2^+^YFP^+^/FOXP2^+^), indicating FOXP2 expression in both glutamatergic and GABAergic neuron populations (**Fig. S1A**). Nearly 80% of YFP⁺ neurons co-expressed FOXP2 (FOXP2^+^YFP^+^/YFP^+^), indicating substantial overlap between FOXP2 expression and Cre-targeted glutamatergic neurons in control mice while in *Foxp2* cKO mice, we confirmed efficient Cre-mediated deletion of *Foxp2* in YFP⁺ neurons (**Fig. 2B**). *Foxp2* cKO FOXP2⁺ cells were also reduced in proportion in *Foxp2* cKO mice (**Fig. 2C,E**), with minimal overlap between FOXP2 and YFP labeling (**Fig. 2B**, **Fig. S1A**). The density of YFP⁺ neurons was also decreased across *Foxp2* cKO ICs (**Fig. 2D**). Residual FOXP2 expression was confined to NEUN⁺ neurons lacking YFP, consistent with retained expression in non-Cre-targeted neuronal populations (**Fig. 2F**).

**Figure 2:**
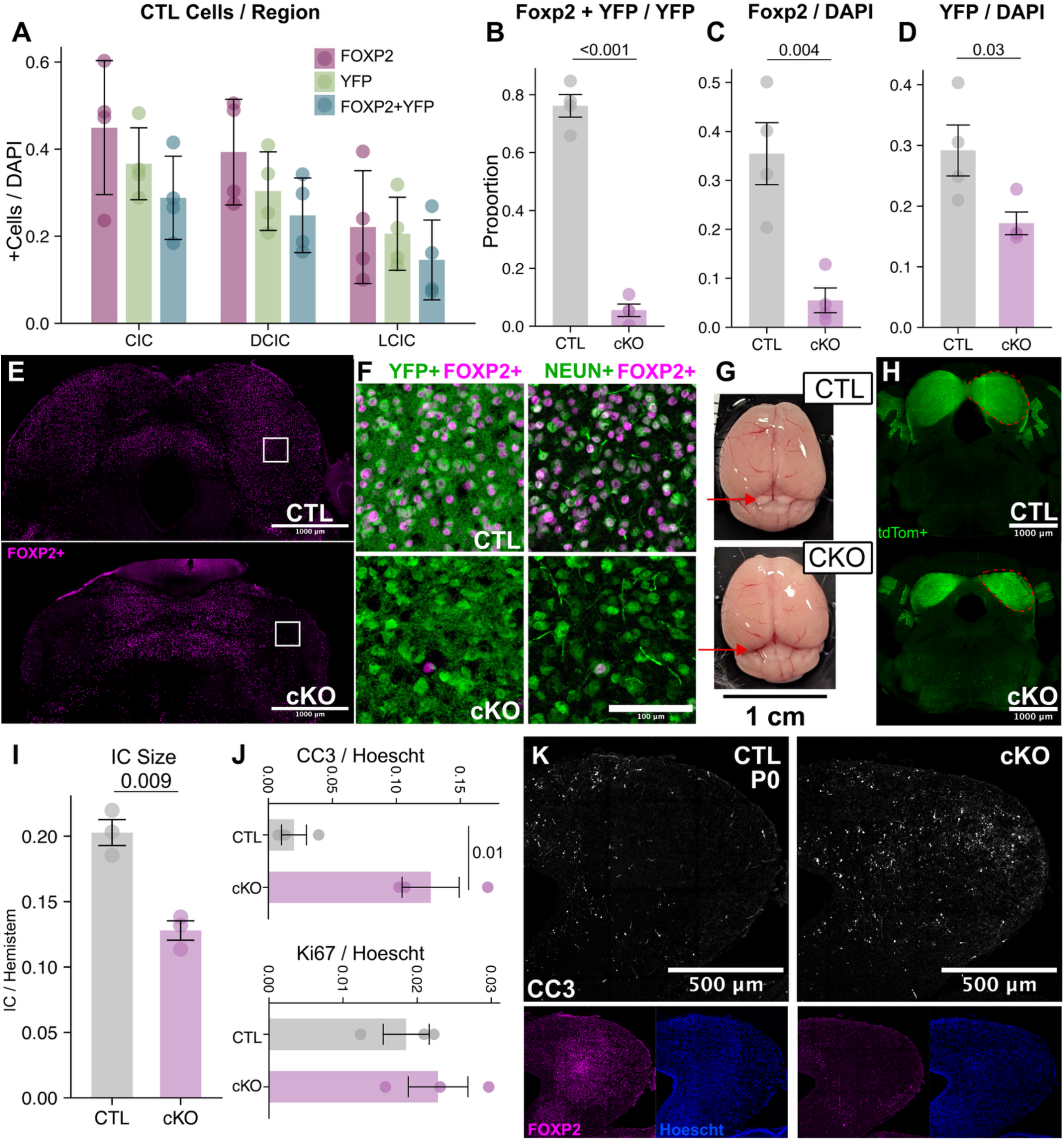
Adult *Foxp2* cKO mice have fewer FOXP2+ cells and smaller ICs with reduced neuronal density. (A) Proportion of cells expressing FOXP2, YFP, and copositive FOXP2 and YFP per region in control (CTL) IC. Error bars are SD. (B) Proportion of YFP+ cells co-expressing FOXP2+ in control and *Foxp2* cKO ICs. Proportion of FOXP2+ (C) and YFP+ cells (D). (E) IHC demonstrates loss of FOXP2 expression and density throughout the IC. White box is approximate area of (F) in the central region of the IC. (F) Higher-magnification view of the central IC. IHC of YFP/Cre reporter (green, left) and NEUN (green, right) with FOXP2 (magenta, both). (G) Coronal sections of control and *Foxp2* cKO brainstem show Cre-activated TdTom reporter signal; red dashed line marks one IC each. (H) Dorsal view of adult control and *Foxp2* cKO freshly dissected brains. Red arrow indicates position of IC. (I) Ratio of Cre-defined IC area to total half-brainstem area. (J) Percent P0 IC cells with CC3+ signal. (K) IHC of coronal sections of the developing dorsal tectum for CC3 (white), FOXP2 (magenta), and Hoescht (blue) for control (left) and *Foxp2* cKO (right) P0 pups. (B-D) Linear mixed-effects models LMM, Error bars are SE; (B: Genotype Effect = 2e^−5^, C: Genotype Effect = 0.036, D: Genotype Effect = 0.0043) (A-D) N = 4 CTL / 4 cKO, ∼3 regions per slice, 1-3 slices per sample. (I) T-test. N = 3 CTL / 3 cKO. (J) LMM. N = 3 CTL/ 3 cKO, 3 regions per sample. Error bars are SE.

Adult *Foxp2* cKO mice gross anatomy showed a smaller IC to the extent that the dorsal cerebellar fold overlapped it, with the IC no longer visible from a dorsal view (**Fig. 2G**). They exhibited a marked reduction in overall IC size, with clear shrinkage evident in coronal sections (**Fig. 2H**). Quantification using serial sections revealed that the average ratio of IC area to the respective brainstem hemisphere area in *Foxp2* cKO mice was approximately half of that observed in control mice (41.5% decrease; **Fig. 2I**).

To determine whether the reduced IC size and reduced number of YFP+ neurons observed in adult *Foxp2* cKO mice reflects altered development or increased death, we examined the timing of *Foxp2* deletion and measured apoptotic cell death during IC development. Previous expression studies in the mouse show *Foxp2* mRNA detectible at embryonic day 16 (E16) in the embryonic dorsal tectum ^5^. Our analysis of the developing dorsal tectum in embryonic tissue revealed that FOXP2 protein is detectable in the IC by E14, while Cre reporter protein appears at E15 and results in efficient loss of FOXP2 expression (**Fig. S1K**). Therefore, our observed changes in the *Foxp2* cKO may indicate that some IC cells expressed *Foxp2* briefly and then lost expression (1-2 days). Other neurons that may have begun *Foxp2* expression post E15 would not have expressed any protein.

To test the hypothesis that increased apoptosis is involved in reducing neuron density in the *Foxp2* cKO ICs, we quantified cleaved caspase-3 (CC3) expression during early postnatal development. At P0, coronal sections of *Foxp2* cKO mice showed an increased proportion of CC3⁺ cells compared to control mice from approximately 2% to 12% indicating increased apoptosis. We also observed no changes in KI67^+^ nuclei proportion, a protein appearing in actively cycling cells, indicating that no compensatory neurogenesis occurred alongside increased apoptosis (**Fig. 2J,K**). These findings suggest that embryonic *Foxp2* deletion followed by elevated apoptosis is involved in the reduced glutamatergic neuron density and the marked reduction in IC size observed in adulthood.

### IC glutamatergic neuron contributions to auditory function

To assess auditory function that may be impacted by the alteration in IC size and neuron density downstream of FOXP2, we performed fundamental auditory behavioral assays that probe IC-dependent modulation of sound processing. The IC contributes to prepulse inhibition (PPI) of the acoustic startle response (ASR), a reflexive subcortical motor response evoked by sudden loud auditory stimuli ^44^. Although generation of the startle response itself does not require the IC, presentation of a preceding non-startle auditory stimulus (a prepulse) engages IC circuitry to suppress the magnitude of the subsequent startle response ^44^. We therefore used ASR and PPI paradigms to evaluate changes in auditory sensitivity and sensory gating with loss of FOXP2.

During the ASR assay, adult mice were exposed to auditory broadband click stimuli ranging from 80 to 120 dB and recorded movement velocity during the reflex response (**Fig. 3A**). Both control and *Foxp2* cKO mice exhibited significantly increased maximum velocity (V-max) movement at 90 dB, relative to their baseline movement recorded in the absence of stimuli, indicating comparable startle thresholds. Startle velocity also remained similar between genotypes at 90 dB and 100 dB intensities but diverged at higher sound levels, with *Foxp2* cKO mice showing significantly increased startle responses at 110 dB and 120 dB (44% and 35% respectively, **Fig. 3B**) consistent with auditory hypersensitivity.

**Figure 3:**
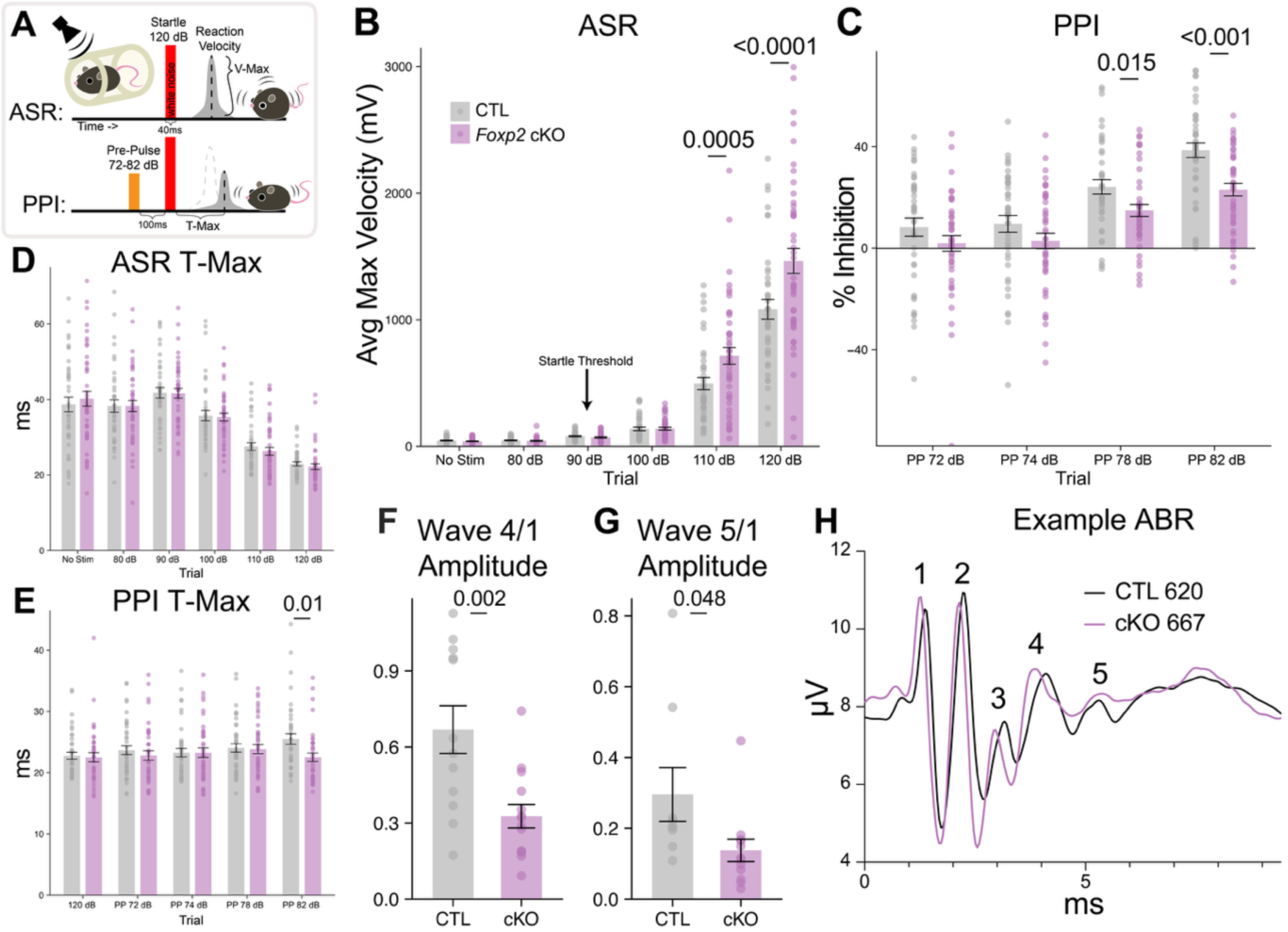
Adult *Foxp2* cKO mice have an increased startle reflex and a deficit in PPI at high pre-pulse, accompanied by weaker ABR wave 4/1 and 5/1 amplitude. (A) Diagram of auditory startle and pre-pulse inhibition behavioral testing. V Max designates the average max velocity recorded post startle stimuli presentation. T-Max is the average time elapsed between stimuli presentation and point of V-Max. (B) Average max velocity at increasing levels of startle stimuli. Stimuli of startle threshold for both genotypes is marked. (C) Percentage of startle inhibition at increasing levels of pre-pulse stimuli. (D, E) Time between startle stimuli and max velocity for no pre-pulse and for 82 dB pre-pulse during the PPI trials. (B-E) LMM with posthoc analysis. (B) Genotype x Trial Interaction effect: p= <0.001. (C) Genotype effect: p= 0.0018. (D) Genotype x Trial Interaction effect: p= 0.9. (E) Genotype x Trial Interaction effect: p= 0.017. N = 41 CTL / 44 cKO. (F) Ratio of amplitude of wave 4 to 1, and (G) wave 5 to 1. (H) Example ABR waves 1 to 5. (F, G) Unpaired t-tests. (F) N= 9 CTL / 8 cKO and (G) N = 6 CTL / 7 cKO. Error bars are SE.

To assess PPI, a non-startling auditory pre-pulse was presented 100 ms prior to a 120 dB startle stimulus. Percent PPI was calculated as the reduction in startle magnitude per trial relative to each animal’s average maximal startle response. Control mice showed increased inhibition of the startle response with increasing pre­pulse intensity. In contrast, *Foxp2* cKO mice exhibited comparably reduced inhibition at higher pre-pulse intensities. During trials with 78 dB or 82 dB pre-pulses, *Foxp2* cKO mice displayed weaker startle inhibition, with an approximately 15% decrease in percent PPI compared to controls during the 82 dB pre-pulse condition (**Fig. 3C**).

The elapsed time between startle stimulus onset and the point of maximal response (T-max) was also measured. During startle-only trials, control and *Foxp2* cKO mice exhibited comparable response speeds from auditory stimulus to peak motor output, with T-max decreasing as response velocity increased alongside startle intensity (**Fig. 3D**). During PPI trials, T-max remained rapid and consistent, comparable to T-max observed in 120 dB startle-only trials. However, in PPI trials with an 82 dB prepulse, *Foxp2* cKO mice showed a reduced T-max compared to controls (post hoc p = 0.1). In contrast, control mice exhibited an increase in T-max at 82 dB relative to their own 120 dB startle-only and 74 dB prepulse trials (p = 0.001 and p = 0.013, respectively) (**Fig. 3E**). This increase in T-max at 82 dB was not observed in *Foxp2* cKO mice.

Since ASR and PPI assays rely on additional brain regions outside the IC, we recorded auditory brainstem responses (ABRs) to compare auditory brainstem region specific signals. To assess auditory brainstem activity across discrete processing stages, we recorded ABRs. Under anesthesia, synchronous neural activity evoked by broadband click stimuli was recorded as field potentials originating from successive auditory brainstem regions. In mice, ABR waves 1 through 5 correspond to activity from the cochlear nerve (wave 1), cochlear nucleus (wave 2), superior olivary complex (wave 3), lateral lemniscus (wave 4), and IC (wave 5 and part of wave 4) (**Fig. 3H**) ^45^. Average ABR wave amplitudes and auditory detection thresholds did not differ significantly between genotypes, nor did peak latencies (**Fig. S2B**). To evaluate IC-associated responses, wave 4 and wave 5 amplitudes were internally normalized to wave 1. *Foxp2* cKO mice exhibited significantly reduced wave 5-to-wave 1 and wave 4-to-wave 1 amplitude ratios compared to control (**Fig. 3F,G**).

Together, these assays reveal altered auditory function in *Foxp2* cKO mice across multiple behavioral and physiological measures. *Foxp2* cKO mice exhibited increased startle magnitude at higher sound intensities, reduced prepulse inhibition at higher prepulse levels, and shorter response latencies during affected PPI trials, despite comparable startle thresholds and baseline response timing. In parallel, auditory brainstem recordings showed preserved early auditory responses but reduced relative amplitudes of IC-associated ABR waves. Collectively, these results indicate that *Foxp2*-dependent alterations of IC glutamatergic neurons modifies auditory sensitivity, sensory gating, and IC-linked brainstem activity.

### Cortical electrophysiological responses to auditory stimuli depend on IC glutamatergic neurons

Since the IC relays processed auditory information to the cortex through the thalamus, we hypothesized that cortical areas could also be affected upon alteration of IC glutamatergic neurons. Therefore, we examined systems-level cortical responses by recording electrocorticography (ECoG/EEG) signals over auditory and frontal cortical areas and measuring auditory event-related potentials (ERPs) (**Fig. 4A**). We first analyzed ERPs evoked by a train of four broadband clicks (**Fig. 4B**, **Fig. S3A-C**). An average ERP response was collected for each mouse, which was then averaged across the genotypic group. We observed clear changes in waveform obtained from both cortical regions in *Foxp2* cKO mice (**Fig. 4**). Five time-epochs were altered based on no overlap between 95% confidence intervals in the average waveform plots (3C-D, marked by colored vertical boxes). For the first stimulus in the train (ERP 1), genotype-dependent changes were observed at later time windows in both auditory and frontal regions (**Fig. 4C**). Quantification of the area under the curve for each nonoverlapping epoch revealed significant differences between *Foxp2* cKO and control mice (**Fig. 4E** (auditory), **3F,G** (frontal)). An earlier ERP 1 component also differed between genotypes at both recording sites (at 97–98 ms for auditory cortex and at 87–106 ms for frontal cortex; **Fig. 4C**, green bar). For the fourth stimulus in the train (ERP 4), a robust genotypic difference was observed only in the frontal cortex at 160–195 ms (**Fig. 4D**). In summary, the large alteration of ERP shape in with loss of FOXP2 is consistent with altered IC function.

**Figure 4.**
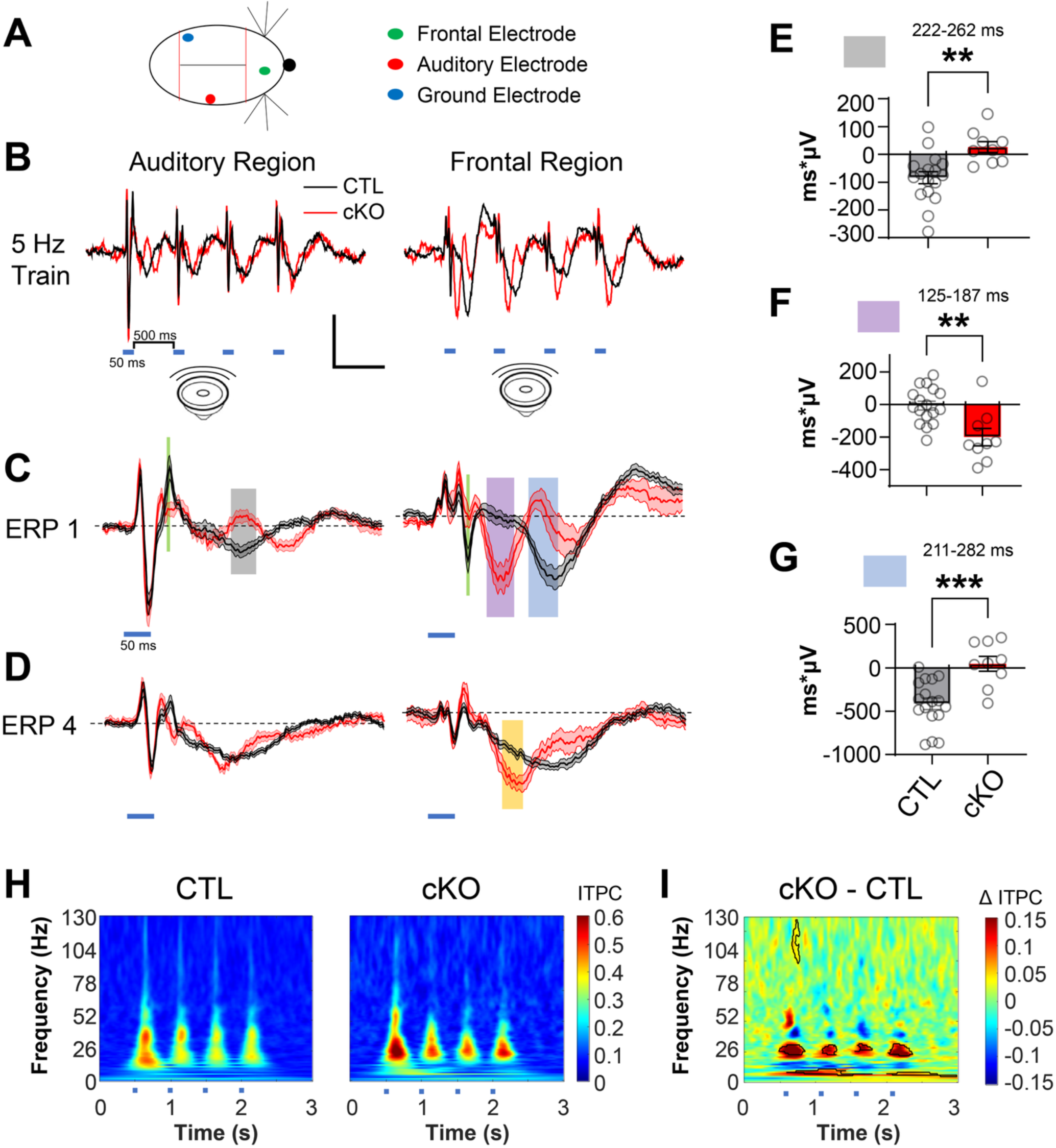
EEG recordings show differences in ERP waveform shape between controls and *Foxp2* cKO mice. A) Diagram of electrocorticography (ECoG) recording locations: ground (blue), auditory electrode (red) and frontal electrode (green). Red lines depict bregma and lambda sutures. B) Average activity measured during the 4 click train (500 ms interval) in control and cKO mice. Blue lines mark the click sounds. C) Average ± SE traces for ERP 1 (response to first click). The colored squares mark time epochs where the 95% confidence intervals of the waveforms are non-overlapping. D) Average ± SE traces for ERP 4. E-G) Areas measured for colored time epochs in C (green and orange not included). Time windows with respect to sound onset are indicated. Error bars are SE. H) Group averages of the time-course of intertrial phase coherence (ITPC) in the auditory region during the sound train (from same data as used in B). Cortical activity is more time-locked to sound stimulation at lower frequencies in the *Foxp2* cKO. Blue lines under axis indicate click stimulus. I) The difference plot of data in H. Statistically significant increases in ITPC (p<0.05) are marked by bold black lines –as observed around 26 Hz. controls are Cre+, flox/WT, and flox/flox mice combined. N = 17 CTL / 9 cKO. P50-60 mice. Scale bars: 500 ms,25 µV for B and 100 ms,25 µV for C-D. ** p<0.01, *** p<0.005.

To assess temporal response properties, we quantified intertrial phase coherence (ITPC) in response to the same auditory click trains. ITPC measures how consistently cortical activity follows the temporal aspects of sound stimuli across trials (**Fig. 4H**). Average difference color plots were generated from ITPC values to compare responses between genotypes (**Fig. 4I**). At the auditory cortical electrode, *Foxp2* cKO mice exhibited higher ITPC values at lower stimulation frequencies (the 4 prominent red patches along the 26 Hz line) compared to control mice.

We further assessed cortical temporal encoding by measuring ITPC using a frequency-modulated “chirp” stimulus. The chirp consisted of a 14 kHz tone modulated by a sinusoid whose frequency increased from 1 to 100 Hz over a 2 second duration (**Fig. S3D**). In average ITPC color plots across genotypes, maximal phase coherence followed the instantaneous modulation frequency of the chirp (**Fig. S3D,E**). Difference plots revealed reduced ITPC in *Foxp2* cKO mice during later portions of the chirp, specifically at modulation frequencies between 70 and 100 Hz (**Fig. S3F**). A similar reduction in ITPC at higher modulation frequencies was also observed at the frontal electrode (**Fig. S3G**). Within the time window highlighted by the white box (SupFig. S3E-G), *Foxp2* cKO mice exhibited reduced peak-to-trough response amplitudes relative to controls, consistent with the decreased ITPC observed during this period (**Fig. S3H**). Together, these results demonstrate that altered IC glutamatergic neurons impair cortical responses to rapidly modulated auditory stimuli.

### Intrinsic excitability of IC glutamatergic neurons is directed by FOXP2

To assess the physiological properties of glutamatergic IC neurons, we performed whole-cell patch-clamp recordings from Cre-dependent YFP-labeled neurons, focusing on the CIC. Intrinsic excitability was assessed by applying incremental depolarizing current steps and quantifying the number of evoked action potentials at each step. Across all current amplitudes tested, *Foxp2* cKO neurons fired fewer action potentials on average than control neurons (**Fig. 5A,B**). Consistent with this finding, these neurons also exhibited decreased input resistance (**Fig. 5C**). We next examined spontaneously occurring excitatory synaptic currents (sEPSCs) where *Foxp2* cKO neurons exhibited reduced sEPSC amplitude compared to control neurons, while sEPSC frequency was unchanged (**Fig. 5D,E,F**). Together, these findings indicate that *Foxp2* loss reduces the intrinsic excitability of glutamatergic IC neurons and weakens excitatory synaptic input strength without altering the frequency of these events.

**Figure 5:**
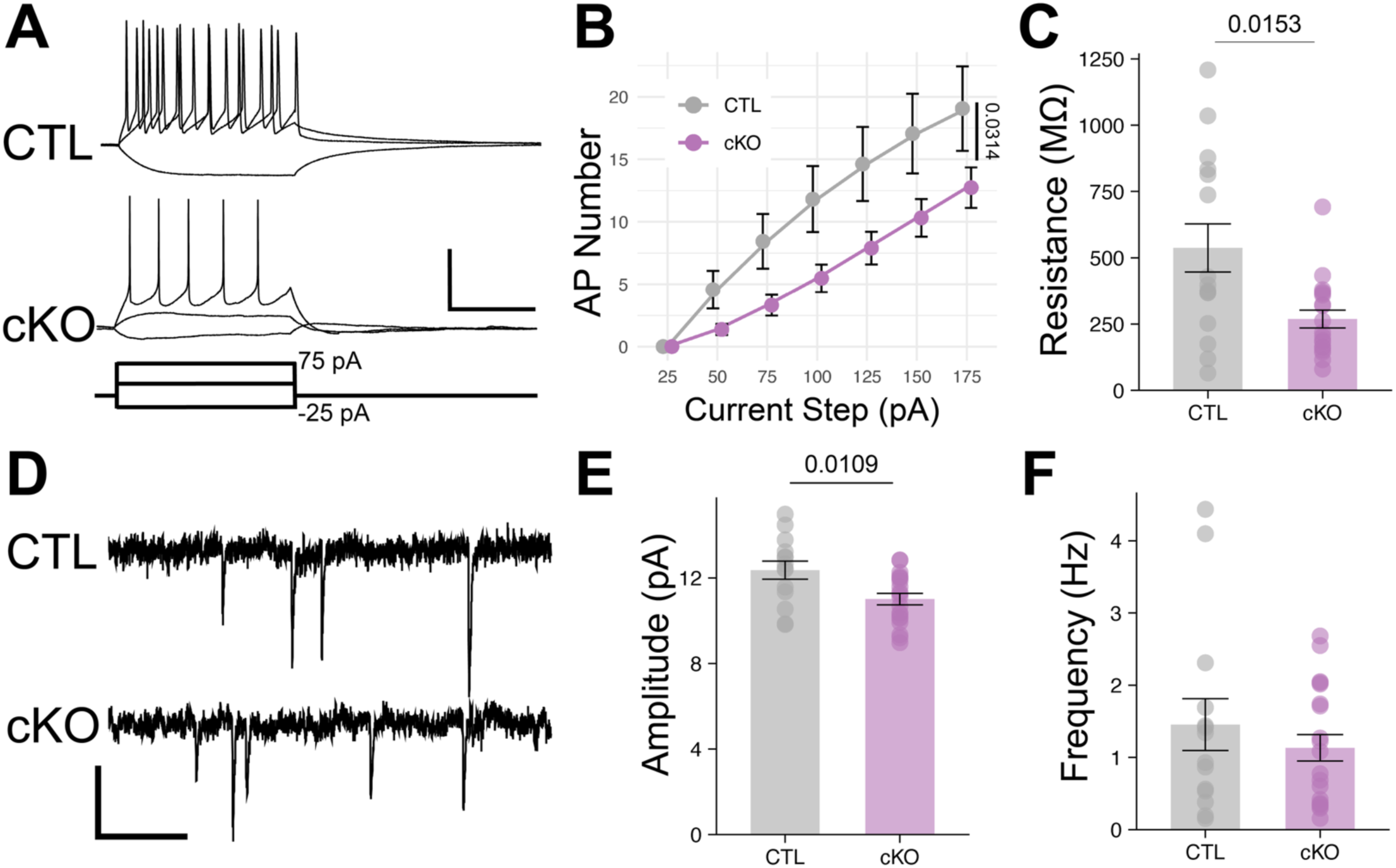
*Foxp2* deletion in Cre-expressing IC neurons of *Foxp2* cKO mice induces hypoexcitability. (A) Traces of whole-cell recordings in Cre-expressing neurons where excitability was measured using current steps. Current steps applied are shown below. (B) Average input-output curves of spike number as a function of step size. The *Foxp2* cKO curve is shifted lower and to the right indicating less excitability relative to control neurons. (C) Input resistance is decreased with *Foxp2* deletion. (D) Traces of spontaneously occurring excitatory synaptic currents (sEPSCs) observed in Cre-expressing neurons of control and *Foxp2* cKO mice. sEPSC amplitude (E) is decreased in *Foxp2* cKO mice while no change is observed in frequency (F). All Cre-expressing cells were identified by a YFP-reporter allele. N = 15 CTL / 20 cKO cells and N = 4 CTL / 6 cKO mice. Control mice are Cre+, YFP+, and Flox-negative. P14-18 mice. Scale bars are: 200 ms, 40 mV for (A) and 200 ms, 10 pA for (D). (B) Mixed-effects analysis, Fixed effects p value<0.0001 for current step, 0.0314 for genotype. (C, E, F) Mann Whitney tests. Error bars are SE.

### Cell subtype-specific proportional changes in the IC

To examine cell subtype-specific contributions to IC-relevant behavior and physiology, we next analyzed multiome data that included IC tissue from the previously profiled controls as well as the newly characterized *Foxp2* cKO mice. Given the reduced size and decreased glutamatergic neuron density observed with loss of FOXP2 (**Fig. 2D,I**), we first examined specific neuronal subtype proportions. We analyzed the combined clustering of control and *Foxp2* cKO samples while applying subtype annotations from control samples to corresponding neuronal clusters in *Foxp2* cKO nuclei. This approach enabled direct comparison of relative population proportions of clusters between genotypes. No novel neuronal cluster identities were detected in the *Foxp2* cKO samples.

Although control samples contained a greater total number of nuclei compared with *Foxp2* cKO samples (**Table S1**), relative population proportions could still be compared. At a broad level, the proportion of excitatory neurons in the dataset was reduced by approximately more than half in *Foxp2* cKO mice relative to controls (**Fig. 6A**). To assess genotype dependent changes in neuronal composition at the subtype level, we analyzed cluster abundances using scCODA with Bayesian compositional modeling and Milo differential abundance analysis as secondary confirmation (**Fig. S5C**). Among excitatory neuronal clusters with high *Foxp2* expression in control samples (>90% of nuclei; **Fig. 1B**), EXC 1, 2, and 3, are significantly underrepresented in *Foxp2* cKO samples. EXC 6, which has moderate *Foxp2* expression, is also decreased. The remaining neuronal clusters show minimal changes in relative abundance (**Fig. 6B**). In control mice, these three *Foxp2*-enriched excitatory clusters together comprise approximately 75% of the excitatory neuronal population but account for only 23% of excitatory nuclei in *Foxp2* cKO samples (**Fig. S5A**). The reduction of nuclei within these clusters is also evident in UMAP visualizations of neuron-only datasets from both genotypes (**Fig. 6C**).

**Figure 6:**
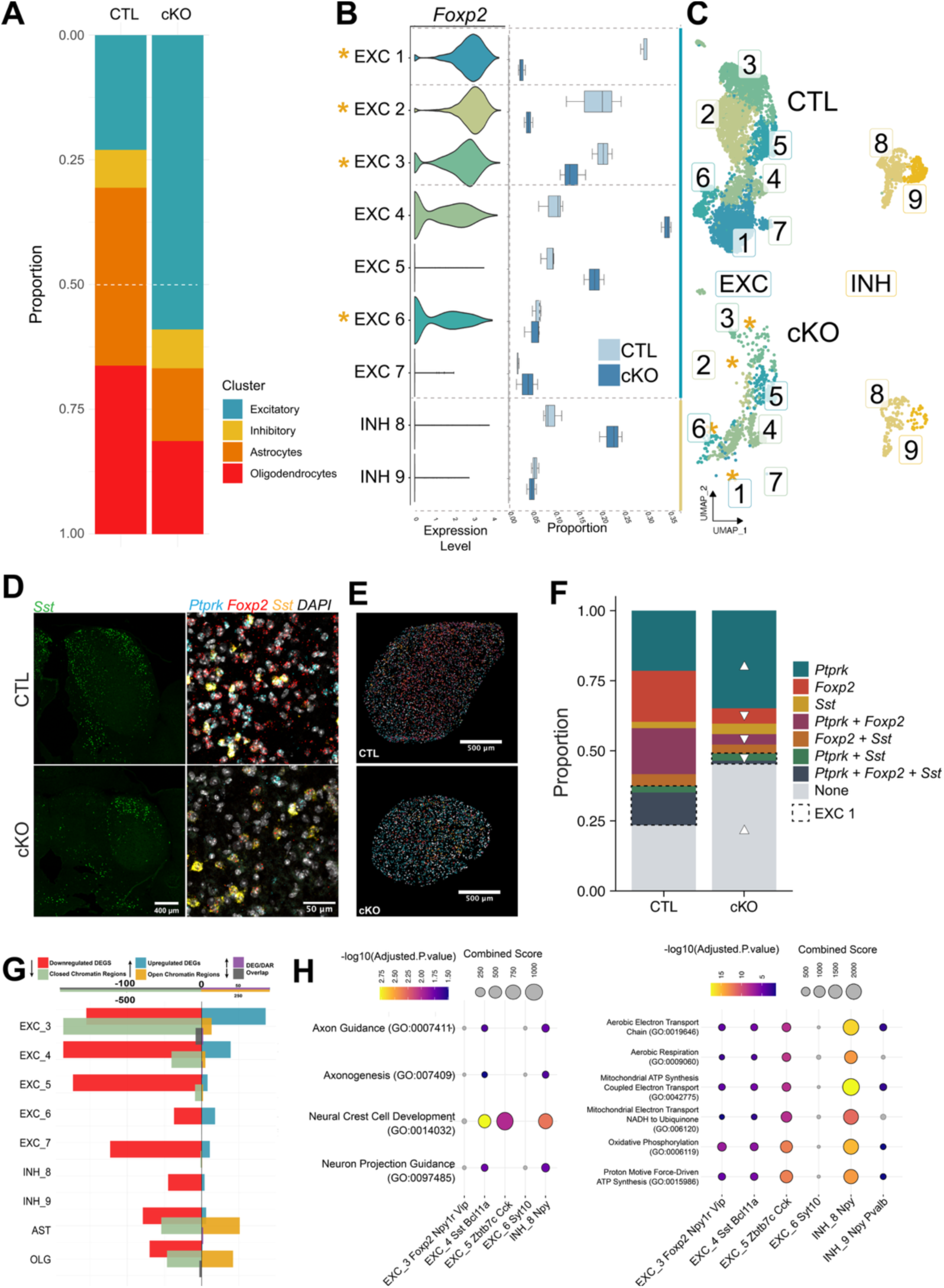
The cellular composition of the IC shows proportional differences between control and *Foxp2* cKO datasets. (A) Bar graphs of proportion of major cell type annotated nuclei in control and *Foxp2* cKO data. (B) Violin plots of normalized expression of marker genes in nuclei by annotated neuronal subclusters (left). scCoda analysis of same neuronal subtypes box and whisker plots of relative neuronal subtype proportions to total nuclei (right). Significantly different clusters are identified by an orange asterisk (*FDR < 0.05). (C) UMAP of sub-setted and re-clustered control (top) and *Foxp2* cKO (bottom) excitatory and inhibitory subtypes. Cluster number labels correspond to subtype annotation. (D) smFISH of control (top) and *Foxp2* cKO (bottom) ICs with probe for Sst (left). Zoomed images in the DCIC of control and *Foxp2* cKO of the dorsal region show *Sst, Foxp2, Ptprk* overlap in control but not *Foxp2* cKO cells (right). (E) DAPI region of interests (ROIs) of nuclei in control (top) and *Foxp2* cKO (bottom) ICs digitally colored by positive probe signal combination. (F) Bar-plots of average proportion of ROIs in ICs marked with each combination of positive probe signals. EXC 1 relevant combinations are outlined in a red dashed box. Chi-squared test of proportions: p < 0.001. Standardized residuals marked with arrowhead if >10. (G) Bar graph of counts of DEGs and DARs per excitatory sub-type. High *Foxp2* expressing clusters are bracketed in red. Downregulated DEGs (red) and closed DARs (green) on the left, upregulated DEGs (blue) and open DARs (yellow) on the right. Downregulated and closed DEG/DAR overlaps (grey) and upregulated and open DEG/DAR overlaps (purple) are also represented. (H) GO analysis (GO Biological Process on top, SynGO on bottom) of upregulated (left) and downregulated (right) DEGs in *Foxp2* clusters EXC 1, 2, and 3.

*Foxp2* expressing affected excitatory clusters can be grouped into two major categories. EXC 2 and EXC 3 are defined by co-expression of *Slc17a6* and *Foxp2*, together with the absence of *Sst*. In contrast, EXC 1 and EXC 4 (an unaffected cluster) are characterized by co-expression of *Foxp2* and *Sst*, distinguishing them from other neurons in which these markers do not co-localize. Within this latter group, *Ptprk* expression further separates the clusters: EXC 1 shows high *Ptprk* expression, whereas EXC 4 lacks *Ptprk*. This distinction between them is relevant, as EXC 1 nuclei are proportionally reduced in *Foxp2* cKO tissue, while EXC 4 nuclei are preserved (**Fig. 6B**, **Fig. S5B**).

To validate the selective loss of excitatory subtypes in intact tissue, we performed single-molecule fluorescent in situ hybridization (smFISH) on coronal sections of adult *Foxp2* cKO IC. Marker combinations derived from the snRNA-seq dataset were used to localize excitatory subpopulations depleted in *Foxp2* cKO samples (**Fig. 6D-F**). In vivo, *Sst*⁺ neurons are predominantly located in the outer cortex of the IC and exhibit characteristic projection patterns ^39^. In control IC, *Sst* expression was prominently localized to the DCIC and LCIC cortical regions, consistent with prior reports (**Fig. 6D**). In *Foxp2* cKO tissue, *Sst*+ cells are also present within the IC; however, gross differences in their spatial distribution are apparent when comparing presumptive dorsolateral and dorsomedial regions. Using combinatorial smFISH labeling for *Foxp2*, *Sst*, and *Ptprk*, we identified cells corresponding to EXC 1 and EXC 4 in both control and *Foxp2* cKO tissue (**Fig. 6D,E**). To account for differences in IC size between genotypes, comparisons were performed using proportions of cells assigned to each molecular identity based on probe expression. Quantification of smFISH signals revealed cells corresponding to the EXC 1 subtype, identified by co-expression of *Foxp2*, *Sst*, and *Ptprk* (including combinations with or without *Sst*-*Ptprk* double positivity), are significantly reduced in proportion in *Foxp2* cKO IC (Chi-squared test p<0.001) (**Fig. 6F**). This decrease mirrors the proportional loss of this glutamatergic subtype observed in the snRNA-seq analysis, despite the overall reduction in IC size in *Foxp2* cKO mice.

### Cell subtype-specific transcriptomic dysregulation in IC glutamatergic neurons

To assess the transcriptional role of *Foxp2* in IC glutamatergic neurons, we performed differential gene expression analysis comparing control and *Foxp2* cKO samples within each IC subtype. This approach enabled evaluation of genotype-dependent gene expression changes within surviving neuronal populations, including cell-autonomous differences in cell types with high *Foxp2* expression as well as non-cell-autonomous changes. To account for unequal numbers of nuclei per sample and avoid cell-level pseudo-replication in clusters affected by proportional loss, we performed differential expression analysis using a pseudobulk approach. Due to the significant loss of *Foxp2* cKO nuclei in the affected clusters, differential gene analysis could not be performed in every cluster. However, EXC 3 and EXC 6, proportionally affected clusters, passed nuclei thresholds and had significant differentially expressed genes (DEGs). EXC 3, a strongly *Foxp2*-expressing cluster, has the greatest number of upregulated DEGs, consistent with FOXP2 as a transcriptional repressor (**Fig. 6G**). In contrast, downregulated DEGs are broadly distributed across neurons and non-neuron cell types.

DEGs from each cluster were assessed for enrichment of functional categories using standard gene ontology (GO) approaches (**Fig. 6H**). Upregulated genes in neuronal clusters are enriched in GO Biological Process terms of neural crest cell development, axonogenesis, and axon guidance. Among SynGO terms, upregulated DEGs belong to pre- and post-synaptic membrane component terms. Downregulated genes across clusters broadly belong to pathways related to aerobic metabolism, specifically oxidative phosphorylation, with varying degrees of enrichment between clusters.

### Juvenile transcriptomic analysis distinguishes developmental functions of *Foxp2*

Because neuronal specification occurs before extensive auditory experience and age, we carried out multiomic experiments in juvenile (∼P14) IC tissue to characterize cell type-specific transcriptional programs spanning both pre- and post-hearing periods and alterations to those programs in the absence of FOXP2. Neurogenesis and initial circuit assembly within the IC occur largely before hearing onset, whereas later post-hearing periods support experience-dependent circuit refinement. P14 corresponds to just after auditory canal opening, enabling comparison of *Foxp2*-dependent gene expression programs before sensory-driven maturation of the auditory system.

Juvenile control and *Foxp2* cKO ICs were processed using the same single-nucleus multiomic workflow as adult samples. Cell types identified in the adult IC were used to annotate juvenile datasets in addition to oligodendrocyte precursor cells, immature oligodendrocytes, and microglia (**Fig. 7A**). Comparing broad cell type proportions between genotypes shows a proportional decrease similar to adult, albeit less dramatic, in the excitatory neuronal proportion (**Fig. 7B**).

**Figure 7:**
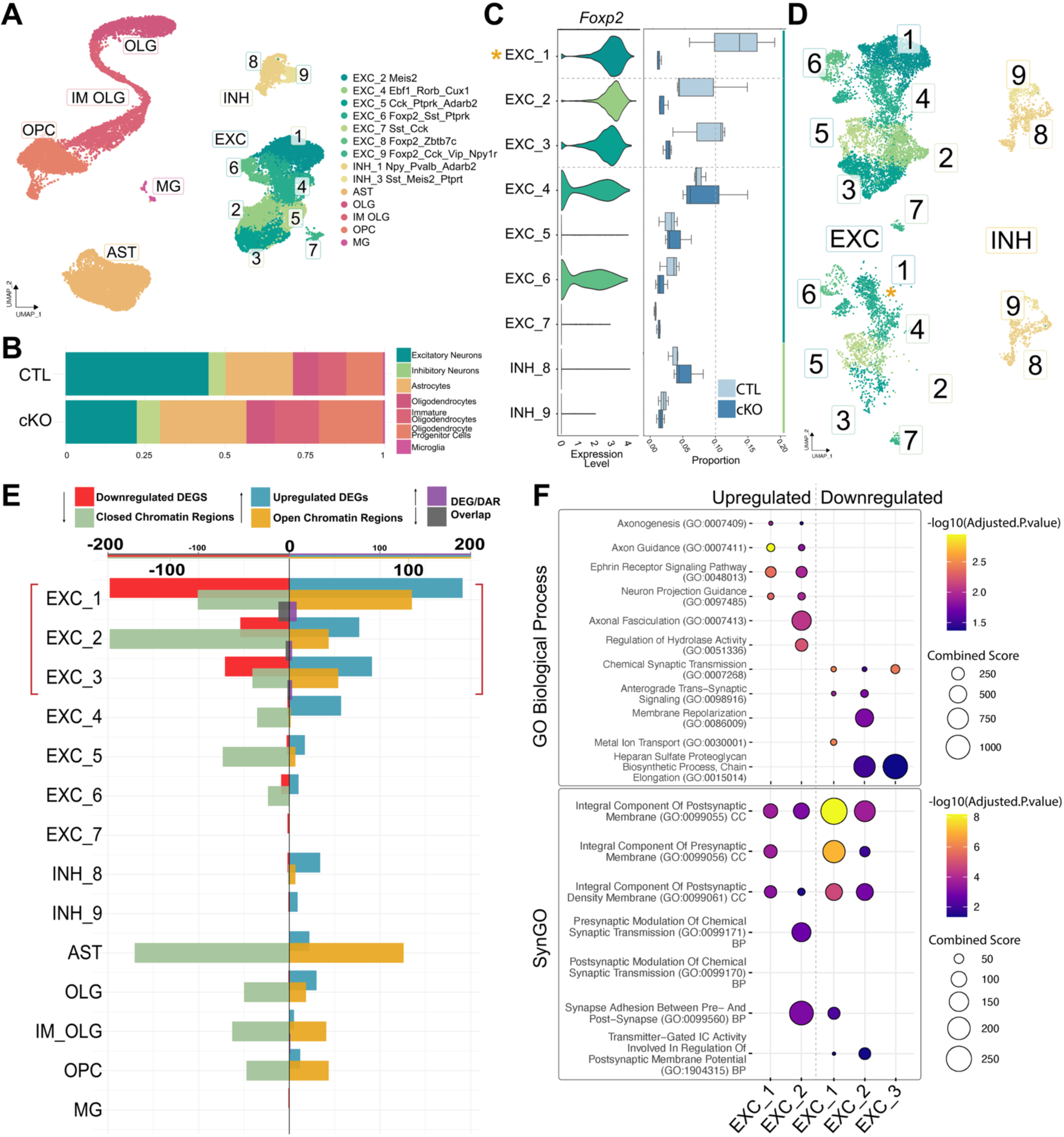
Juvenile control and *Foxp2* cKO IC multiome recapitulate only some adult neuron subtype proportional differences with detectable DEGs. (A) UMAP of juvenile control nuclei labeled with transferred neuronal annotation. (B) Proportion of broad cell types in juvenile nuclei data within control and *Foxp2* cKO datasets. N = 3 control samples / 3 cKO samples, each sample is pooled from 2 individuals of the same sex, age range, and genotype. (C) Violin plot of *Foxp2* expression per neuronal subcluster (left). scCODA proportional differences between genotypes (right), asterisks indicate significant proportional abundances. (D) UMAP of sub-setted and re-clustered control (top) and *Foxp2* cKO (bottom) excitatory and inhibitory subtypes. Cluster number labels correspond to subtype annotation. (E) Bar graph of counts of DEGs and DARs per excitatory sub-type. High *Foxp2* expressing clusters are bracketed in red. Downregulated DEGs (red) and closed DARs (green) on the left, upregulated DEGs (blue) and open DARs (yellow) on the right. Downregulated and closed DEG/DAR overlaps (grey) and upregulated and open DEG/DAR overlaps (purple) are also represented. (F) GO analysis (GO Biological Process on top, SynGO on bottom) of upregulated (left) and downregulated (right) DEGs in *Foxp2* clusters EXC 1, 2, and 3.

Marker gene expression patterns in juvenile control samples closely mirror those of corresponding adult subtypes, and *Foxp2* expression remains highest in excitatory clusters EXC 1, 2, and 3 (**Fig. 7C**). Cell type proportions were assessed using scCODA. Unlike adult tissue, juvenile *Foxp2* cKO samples exhibits underrepresentation of only excitatory cluster EXC 1, whereas EXC 2, 3, and 6 proportions are not significantly reduced at this developmental stage. No other neuronal clusters showed significant overrepresentation in juvenile *Foxp2* cKO samples (**Fig. 7C,D**).

Differential gene expression and chromatin accessibility analyses were performed on juvenile datasets using the same methods applied to adult samples. In the juvenile samples, clusters EXC 1, 2, and 3 all had sufficient nuclei within both control and *Foxp2* cKO samples to perform DEG analysis. As in adult tissue, the majority of DEGs and DARs are concentrated within *Foxp2*-enriched excitatory clusters (**Fig 6E**). However, juvenile *Foxp2* cKO samples did not exhibit widespread downregulation of metabolic genes observed in adult clusters.

GO analysis of juvenile downregulated DEGs of *Foxp2* clusters revealed strong enrichment for synaptic and ion channel–related terms (**Fig 6F**). Neurodevelopmental and axonal targeting pathways are enriched in downregulated DEGs. Among the most strongly dysregulated genes, we identified multiple axonal targeting molecules whose proteins function as receptors or ligands that guide extending axons via local attractive and repulsive cues. While EXC 3 exhibited similar transcriptional patterns, significant enrichment of these upregulated axon guidance genes was restricted to EXC 1 and EXC 2.

Notably, members of the Eph receptor and ephrin ligand families are among the dysregulated genes shared across all three *Foxp2*-enriched excitatory clusters (**Fig. 8A**). These genes are known to participate broadly in axon targeting as well as additional developmental processes including proliferation, differentiation, migration, and synaptogenesis, depending on expression location and developmental stage ^46^. As an example of cluster-specific transcriptional dysregulation, differential expression analysis of EXC 1 identified *Epha4* as one of the most significantly upregulated genes in *Foxp2* cKO neurons compared to control (**Fig. 8A**). In the developing IC, *Epha4* expression is patterned, with a gradient along the tonotopic axis of the CIC and localization to discrete modular structures within the LCIC ^47^. In contrast, EXC 1 neurons represent a large population of glutamatergic *Sst*⁺ neurons that lacks strong *Epha4* expression and are positioned dispersed around these canonical *Epha4*-enriched domains. The observed upregulation of *Epha4* in EXC 1 neurons therefore reflects altered expression of an axon guidance receptor in a neuronal population not normally associated with this molecular pattern.

**Figure 8:**
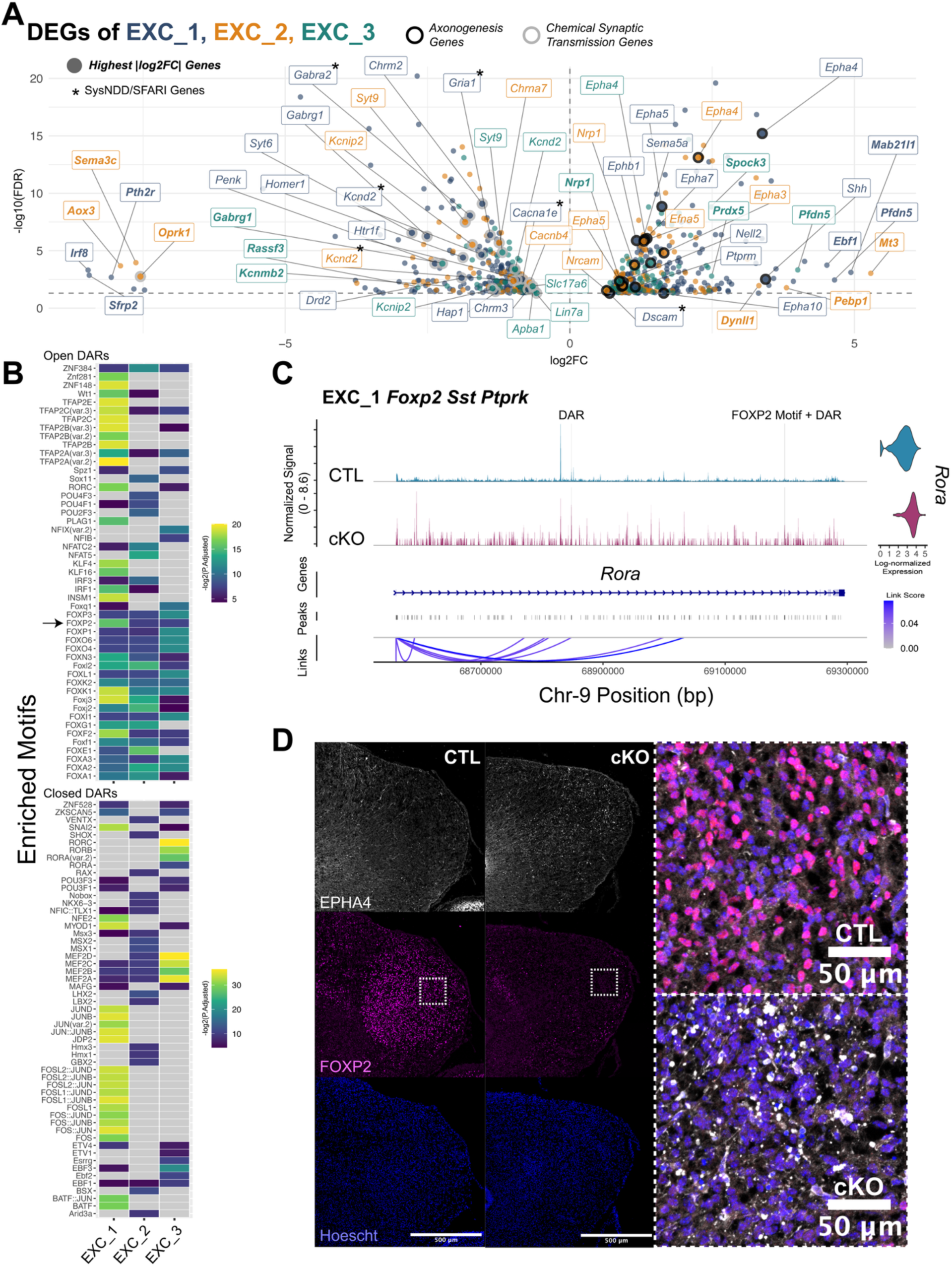
Comparison of control and *Foxp2* cKO cell subclusters reveals a graded molecular response that correlates with *Foxp2* expression level in the control condition. (A) Combined volcano plot of significant DEGs within EXC 6, 8, and 9 colored by cluster. Known axonogenesis genes (bolded dot outline) and the top log_2_FC genes (bolded names) are labeled. Genes marked with an asterisk are included in the SysNDD Definitive and/or SFARI Score 1 or Syndromic gene database. (B) Heatmap of enriched motif accessibility colored by -log10(adjusted p value) and clustered by similarity of expression between cell types. (C) Coverage plot of ATAC-seq derived peak signals within the *Epha4* gene DNA location between genotypes in EXC 6. DAR peak and DAR peak with FOXP2 motif location are highlighted (grey). Bottom links showing relationships between genomic positions and the 3’ region. Top right is normalized gene expression. (D) IHC of P0 ICs (left) showing *Foxp2* cKO *Foxp2* (magenta) absent and *Epha4* (white) increased expression in the developing IC. (right) Zoomed in dashed square insert shows a composite image.

To determine chromatin-level regulatory changes associated with *Foxp2* loss, we analyzed paired snATAC-seq data from control and *Foxp2* cKO samples. Differential chromatin accessibility was assessed using a pseudobulk strategy with the same quality control parameters applied to the transcriptomic analysis. The greatest number of differentially accessible regions (DARs) within neural populations is observed in excitatory clusters with EXC 1, EXC 2, and EXC 3, mirroring the cluster-specific patterns observed in differential gene expression (**Fig. 7A**). We next linked the nearest gene to each DAR for GO analysis, however, no significant GO terms were enriched in the juvenile DAR data.

To assess the regulatory relevance of DARs, we identified transcription factor motif enrichment within the DARs (**Fig. 8B**). EXC 1, 2, and 3 showed significant enrichment for FOX family motifs, including the FOXP2 motif, within open DARs. In addition, EXC 1 exhibits enriched motifs corresponding to FOS, JUN, and TFAP family transcription factors within closed DARs. We also found enrichment of MEF2 family motifs in closed DARs in EXC 1, 2, and 3, including MEF2C, consistent with other reports in the striatum ^29^.

To examine potential regulatory relationships between chromatin accessibility and gene expression, we assessed overlap between DEGs and DARs. EXC 1 exhibited the greatest number of overlapping DEGs and DARs, consistent with its high degree of transcriptional and chromatin dysregulation, with 20 genes shared (Fisher’s exact test FDR = 0.0008 upregulated, 6.51E^−10^ downregulated). These overlapping genes included *Rora*, which encodes a nuclear receptor protein with associations to NDDs and is a syndromic identified gene in the SFARI database ^48^. *Zmiz1* is another upregulated and open chromatin-associated gene that has roles in dendritic and synaptic development and is associated with NDDs and ASD ^49^. In *Foxp2* cKO EXC 1 neurons, loss of *Foxp2* was associated with increased chromatin accessibility near *Rora* and *Zmiz1* accompanied with increased transcript abundance. We identified a FOXP2 motif within a DAR in the *Rora* locus in EXC 1 neurons (**Fig. 8C**). These findings indicate that loss of *Foxp2* is associated with coordinated changes in chromatin accessibility and gene expression at loci involved in axonogenesis and related developmental programs.

### Altered EPHA4 protein patterning in early postnatal *Foxp2* cKO IC

To assess whether transcriptomic and chromatin changes were reflected at the protein level, we examined EPHA4 expression during early postnatal development. *Epha4* was selected based on its strong upregulation in *Foxp2* cKO neurons and its established role in axonal targeting. Given that EPHA4 expression is highest during developmental periods of axon targeting and refinement, IHC for EPHA4 was performed on P0 brains (**Fig. 8D**). In *Foxp2* cKO mice, developing ICs exhibits increased density of EPHA4-positive puncta compared to controls, with ectopic expression observed in both central and cortical regions of the IC. Although cell type specific expression could not be resolved at the protein level, these findings demonstrate altered spatial patterning of EPHA4 in the developing IC following loss of *Foxp2*.

### Metabolic gene expression diverges between juvenile and adult *Foxp2* cKO neurons

To compare metabolic gene regulation across developmental stages, juvenile and adult datasets were integrated and jointly normalized. We examined expression of nuclear-encoded oxidative phosphorylation genes to determine whether observed metabolic changes were age-dependent or genotype-dependent. Clustering of oxidative phosphorylation genes, including those identified in adult DEGs, by genotype and age revealed a clear separation between adult control and adult *Foxp2* cKO samples (**Fig. S5D**). Relative to juvenile levels, adult *Foxp2* cKO neurons failed to upregulate oxidative phosphorylation genes to control levels and, in some cases, exhibited reduced expression compared to juvenile tissue. Analysis of the integrated dataset therefore revealed divergent regulation of metabolic gene expression between juvenile and adult stages in control versus *Foxp2* cKO neurons.

## Discussion

### IC cell types and FOXP2 populations

IC cell subtype identification has previously relied on non-transcriptomic classifications with limited success in identifying consistent cell identities. Single-nucleus transcriptomic approaches now provide an opportunity to integrate previously separate observations with molecular definitions of neuronal identity via gene expression patterns, enabling more direct links between IC cell types and function. One transcriptomically defined cell type is the *Foxp2*+*Sst* co-expressing glutamatergic population, which represents a major subset of *Sst*-expressing neurons primarily located in the DCIC and LCIC regions. Prior work has shown that IC *Sst* neurons mainly project to secondary auditory thalamic regions (notably the posterior limiting nucleus), suggesting a role in secondary auditory processing based on both IC localization and projection targets ^30,36^. The specific spatial and projection organization of *Foxp2*-expressing *Sst* glutamatergic neurons within this broader population remains unresolved, but evidence suggests they contribute to multi-sensory integrative processing within IC circuits, with extra-auditory and cortical region inputs. The enrichment of FOXP2 within these defined glutamatergic populations raises the possibility that FOXP2 is required for their function. We note that a recent paper profiled the IC in adult wildtype mice and similarly defined neuronal subtypes ^51^. However, this study did not delineate *Foxp2*-expressing cell types. Thus, a future study will need to determine the correspondence and overlapping annotations between these datasets. The major disruption in IC neuron proportions induced by *Foxp2* deletion suggests that FOXP2 plays a critical role in maintaining the integrity of specific glutamatergic neuron populations in the IC during development. FOXP2 may regulate developmental identity programs necessary for the integration of IC neurons into maturing auditory circuits, potentially through regulation of axon guidance and synaptic development programs.

### Developmental circuit assembly mechanisms

Axon guidance programs provide a plausible developmental mechanism linking early FOXP2-dependent gene regulation to long-term circuit dysfunction in the IC. Cell adhesion, neurite outgrowth, and axonogenesis are known roles of targets under FOXP2 regulation across regions and mutations. Because axonal targeting and arbor refinement occur primarily during late embryonic and early postnatal development, dysregulation of guidance signaling and synaptic structure during this period could produce lasting alterations in circuit organization ^6^.

Eph/ephrin signaling is a key regulator of spatial patterning in the developing IC, particularly in establishing tonotopic organization of incoming auditory projections ^46,47^. Within the differentially expressed genes of FOXP2-enriched clusters, most notably EXC 1 (*Foxp2 Sst Ptprk*), we identified upregulation of *Epha4*, among other members of the Eph/ephrin families. Although gene dysregulation is readily detectable in the juvenile IC, the functional consequences of altered guidance signaling are likely to arise earlier during developmental windows when axonal targeting and refinement are actively occurring. During the peak period of axonal patterning around birth, we observed increased and ectopic EPHA4 protein expression in nodular structures throughout the IC, suggesting that *Foxp2* loss alters spatial guidance cues during the window when incoming projections are being organized

### Synaptic Regulation Under FOXP2

Integrated chromatin accessibility and motif enrichment analyses revealed enrichment of FOX family transcription factor motifs, including FOXP2 itself, within differentially accessible regions. While adult DARs were limited, we found open and closed DARs mostly limited to EXC 1, 2, and 3 in neuronal clusters of the juvenile data. Consistent with these regulatory changes, transcriptional analysis revealed dysregulation of genes controlling synaptic organization and neuronal excitability. Notably, potassium channel genes including *Grik1* and *Grik4*, which are upregulated in *Foxp2* cKO EXC 1 neurons, are kainite-type glutamate receptors, ligand-gated ion channels involved in synaptic plasticity and NDDs such as ASD and schizophrenia ^52–55^. *Kcnj6* (GIRK2, potassium inwardly rectifying channel) ^56,57^, *Kcnq5* (M-type, voltage-gated potassium channel subunit) ^58,59^, and *Kcnd2* (A-type, voltage-gated potassium channel) ^58,60^, which are downregulated in *Foxp2* cKO EXC 1 neurons, are regulators of membrane excitability and neuronal responsiveness and have been implicated in NDDs and epilepsy.

Metabotropic glutamate receptor genes, including *Grm1* and *Grm8*, are also dysregulated. *Grm8*, in particular, is upregulated across multiple FOXP2-expressing glutamatergic subtypes, suggesting a shift in presynaptic modulation of glutamatergic signaling^61,62^. In parallel, *Sorcs3* and *Sorcs1*, which encode transmembrane receptors involved in glutamate receptor trafficking and synaptic organization, are similarly upregulated, indicating altered postsynaptic receptor dynamics and plasticity ^63,64^. *Sorcs3* and *Foxp2* together have been identified as genes likely to be relevant to ASD and ADHD ^12,65^.

### Cellular and Circuit Physiology

Consistent with dysregulation of synaptic and axon guidance pathways and selective loss of excitatory neurons, *Foxp2* cKO mice exhibited alterations across multiple levels of auditory processing. Because the IC contributes inhibitory modulation within startle circuitry^44^, disruption of IC excitatory populations may shift excitation–inhibition balance within auditory gating networks, resulting in the observed auditory hypersensitivity and impaired sensory gating. The relatively modest magnitude of behavioral disruption despite substantial anatomical and molecular changes suggests either the limited involvement of the affected cells in ASR and PPI or the presence of compensatory mechanisms within the auditory brainstem or midbrain, potentially involving altered gain control across IC feedback circuits.

Consistent with impaired IC function, ABR recordings revealed diminished neural output at waves associated with IC activity, while earlier brainstem waves remained intact. Previous work with FOXP2 mutations reported either normal ABR with a heterozygous loss-of-function mutation (S321X) or prolonged latencies and reduced wave amplitudes with a dominant-negative mutation (R552H)^45^. Although reduced IC-associated ABR amplitudes appear inconsistent with heightened startle reactivity, these assays probe distinct aspects of auditory processing. ABR reflects synchronous activity across the ascending auditory pathway, whereas ASR engages specialized reflex circuits linking cochlear nucleus outputs to brainstem motor pathways. Disruption of IC-dependent modulation of these circuits may therefore amplify behavioral responses without increasing early auditory brainstem activity.

Changes in IC function were also reflected in downstream cortical responses, which suggested impaired temporal coordination of cortical responses to repeated sounds. Because the IC provides the principal subcortical relay to auditory cortices via the thalamus, disruption of IC computations can propagate through ascending pathways and alter higher-order sensory representations^46^. At the cellular level, surviving glutamatergic neurons in the CIC showed reduced intrinsic excitability, a change that would be expected to weaken coordinated IC output and contribute to the reduced IC-associated ABR amplitudes observed in vivo.

*Foxp2*-expressing glutamatergic neuron subtypes make up approximately three-quarters of the glutamatergic neuron population, and their loss is therefore likely to alter excitation–inhibition balance within IC circuits, necessitating compensatory adaptations in remaining neurons that may impair more complex forms of auditory processing. Moreover, disruption of excitatory subtypes associated with *Sst*- and *Vip*-linked projection pathways to thalamic targets may limit the neurons responsible for information relay, effectively creating a bottleneck in dimension-rich auditory circuit signaling ^39^. The regional and functional consequences of each subtype will be important for future work to follow up on.

### Network-level metabolic adaptation

The widespread downregulation of oxidative metabolism genes across cell types suggests a largely non–cell-autonomous disruption of metabolic programs in the adult *Foxp2* cKO IC. During normal development, neurons in the auditory system undergo a postnatal increase in oxidative metabolism following the onset of auditory experience (around 2–3 weeks of age in mice), reflecting the rising energetic demands of circuit maturation and sustained synaptic activity ^47^. Given the high baseline metabolic demand of the IC and the reliance of mature neurons on oxidative phosphorylation ^48^, this pattern is therefore unlikely to result solely from direct FOXP2 regulation and instead likely reflects secondary consequences of disrupted circuit organization and sustained network stress following early developmental perturbations. One possible explanation for this metabolic dysregulation is an excitation–inhibition imbalance emerging during the onset of sensory-driven activity following early loss of excitatory neurons. Such a mismatch between network activity and metabolic capacity could impair neuronal maturation and reduce the functional resilience of IC circuits.

### Disease relevance of FOXP2-dependent IC dysfunction

FOXP2 has been strongly implicated in NDDs, including childhood apraxia of speech, ASD, ADHD, and schizophrenia, raising the possibility that disruption of FOXP2-dependent transcriptional programs in the IC may contribute to altered auditory processing relevant to these conditions. For example, *Rora*, which we observed to be both upregulated and more accessible in EXC 1 *Foxp2 Sst Ptprk* neurons, encodes a retinoic-acid-receptor-related orphan receptor-α strongly connected with ASD ^48,71,72^, whose transcriptional targets overlap with FOXP2 (*NTRK* and *A2BP1*) ^73^.

The coordinated increase in chromatin accessibility and transcriptional upregulation at this locus therefore supports a conserved repressive role for FOXP2 in regulating transcription factors, synaptic, and circuit-stabilizing genes within IC excitatory neurons. Consistent with this interpretation, *Foxp2* loss also disrupted the expression of additional genes involved in synaptic adhesion and circuit stability, including *Cntnap4*, *Meis2*, *Cdh9*, and *Dscam,* all of which have strong genetic associations with NDDs ^74–78^. In addition to synaptic genes, we show that FOXP2-dependent transcriptional programs in the IC also regulate axon guidance pathways, including Eph/ephrin signaling, suggesting that FOXP2 loss may disrupt both the assembly and stabilization of auditory circuits.

### Limitations of the Study

The Nstr1-Cre mouse is specific for a subpopulation of excitatory neurons in the IC, but it is also expressed in the olfactory bulb, the striatum, parts of the cerebellum, and limbic cortex ^79,80^. Therefore, it is possible that altered behavior and EEG in the *Foxp2* cKO mice resulted from *Foxp2* in these areas and not in the IC. Three factors mitigate this possibility. First, Cre was expressed in many cells that do not express *Foxp2*. Therefore, those cells and regions have no or minimal loss-of-function of *Foxp2*. Second, the regions and cell types that do undergo loss-of-function for *Foxp2* are not principal regions of auditory processing and *Foxp2* deletion in these areas is unlikely to affect our systems-level data, as our focus is on simple auditory processing, both EEG and behavior. And third, among all the auditory structures, the IC had the greatest overlap and loss-of-function for *Foxp2*. For example, within the auditory system, Cre is also expressed in the cochlear nucleus, but this region does not show FOXP2 coexpression in our stains (**Fig. S1B,E**) and thus does not contribute to FOXP2 loss-of-function. Among auditory structures, Cre and FOXP2 are most strongly co-expressed in IC glutamatergic neurons, resulting in the regional specificity for FOXP2 loss-of-function to the IC when focusing on auditory system changes.

Other limitations of the present study point to critical directions for future work. Although our data demonstrate that FOXP2 directs IC cell type composition, gene regulation, and auditory-related behavior, we did not directly assess how these changes affect context-dependent sound processing or stimulus-specific coding within the IC. This limitation arises from the complexity of IC circuitry, which integrates diverse ascending, descending, and commissural inputs in a highly cell type–dependent manner. As a result, while we observe behavioral and molecular disruptions, we cannot yet determine how *Foxp2*-dependent changes alter specific circuit computations or output channels. In particular, we did not directly examine whether altered proportions or transcriptional states of IC neuron subtypes correspond to changes in synaptic connectivity, projection targets, or tonotopic organization. Without circuit-level tracing or physiological mapping, it remains unclear whether the observed molecular changes primarily affect internal IC processing, long-range output routing, or both. Similarly, while auditory startle and related assays indicate functional consequences of *Foxp2* deletion, these paradigms do not resolve how spectral tuning, frequency discrimination, or adaptive auditory learning are impacted. Consequently, our findings establish FOXP2 as a regulator of IC neuronal identity and integrity but stop short of defining its role in shaping precise auditory representations or higher-order sensory computations.

## Conclusion

Overall, these findings detail transcriptional programs that coordinate the developmental assembly and long-term stability of glutamatergic circuits in the IC. Disruption of these programs alters axon guidance, synaptic organization, and intrinsic neuronal excitability, ultimately impairing IC circuit output and downstream auditory processing. The widespread metabolic dysregulation observed across cell types further suggests that compensation for these circuit disruptions imposes sustained energetic demands on IC networks. These results therefore identify the IC as a critical subcortical locus through which FOXP2-dependent developmental programs shape auditory circuit function. More broadly, our findings suggest that sensory and behavioral phenotypes associated with neurodevelopmental disorders may arise not only from cortical dysfunction but also from early disruptions in subcortical sensory circuits that influence downstream neural activity and behavior.

## Methods

### Mice

Mice were housed in an accredited vivarium on a 12-h light/dark cycle. Food and water were provided ad libitum. Genotypes were determined using PCR analysis of genomic DNA isolated from toe or tail clippings. All procedures were approved by the University of Texas Southwestern Medical Center Institutional Animal Care and Use Committee. Experiments were conducted in accordance with the NIH Guide for the Care and Use of Laboratory Animals. Mice of both sexes were used for all experiments.

Mice used for this paper are *Foxp2*^flox/flox^ (French et al. 2007; #026259, Jackson Laboratory), Ntsr1-cre GN209 (GENSAT BAC transgenic), R26R-EYFP (#006148, Jackson Laboratory), Ai14 (#007914, Jackson Laboratory). Mice were maintained on a C57bl/6 J background for at least 8 generations when first obtained by the lab, and after that, maintained by interbreeding with WT C57bl/6 J mice (Jackson Laboratories).

We performed *Foxp2* conditional deletion experiments by crossing the Ntsr1-Cre mouse (GN209) ^1–4^ with the *Foxp2*^flox/flox^ mouse ^5^ to produce *Foxp2* cKO mice. Even though the Ntsr1-Cre line expresses Cre in non-auditory structures outside of the IC, it is still sufficient for targeting the IC when limiting experiments to the auditory system. *Foxp2* is not expressed in all cells, and therefore, in those cells, there is no change in *Foxp2* expression. As a result, only cells that express both Cre and *Foxp2* experience loss in FOXP2, which in turn, increases the specificity of the cKO strategy for targeting the IC (summarized in **Fig. S1**). The Ntsr1-Cre mouse expresses Cre most strongly in the IC (in all 3 subregions) and in a small number of cochlear nucleus neurons in the CN and CRN, and all are likely glutamatergic ^3,6–9^. Our recordings of Cre+ cells in the IC indicate they all fire action potentials (20/20 cells), and therefore, are indeed, neurons. There is Cre-expression in brainstem structures not directly related to auditory processing (including the stratum opticum of the superior colliculus ^1,2^) and in many axon fibers, which we believe unlikely to significantly impact our results. Cre is expressed before E15 (**Fig. S1K**) and results in Cre-dependent reporter expression by E15 ^3^. To our knowledge, there is no report for *Ntsr1* expression in IC neurons suggesting that Cre is ectopically expressed ^10^.

Behavior, snRNA, and snATAC-seq experimental mice (*Foxp2* cKO) and control littermates were generated by crossing *Foxp2*^flox/flox^ with Ntsr1-cre^+/wt^:*Foxp2*^flox/flox^. Imaging experimental and control animals were generated by crossing *Foxp2*^flox/wt^ with Ntsr1-cre^+/wt^: *Foxp2*^flox/wt^: R26R-EYFP^+/+^ or Ai14 tdTomato^+/+^ cre reporter lines to generate cKO *Foxp2*^flox/flox^: Ntsr1-cre^+/wt^: R26R-EYFP^+/wt^ or Ai14^+/wt^ and control *Foxp2*^wt/wt^: Ntsr1-cre^+/wt^: R26R-EYFP^+/wt^ or Ai14^+/wt^ littermates.

### Timed Breeding

To accurately capture embryonic stages of development, dams were vaginally swabbed for estrus stage before being paired with a singly-housed male overnight. The dam was then separated and monitored for weight gain over 1-2 weeks. Separation day is considered embryonic day 0.5 (E0.5).

### Immunohistochemistry

Adult and juvenile mice were anesthetized by injection with 80-100mg/kg Euthasol and transcardially perfused with 4% PFA. Neonatal P0 pups were euthanized with rapid decapitation into ice-cold PBS. Brains were then post-fixed overnight in 4% PFA and then cryoprotected in 30% sucrose in PBS for embedding in Tissue-Tek CRYO-OCT Compound (catalog #14-373-65, Thermo Fisher Scientific). Cryosectioned tissue was sliced at 20-40 um and stored free-floating in 1X PBS solution or directly to glass slides. Staining was performed on free-floating sections or on glass slides.

Embryonic mice were harvested from a pregnant dam following CO_2_ euthanasia. The dissected uterus was placed in ice-cold PBS to isolate each pup from the organ. After collecting a tail DNA sample, the pup was decapitated and head placed in 4% PFA for 4-12 hours depending on age. After cryoprotection and embedding, tissue was cryosectioned at 20-30 um directly to glass slides. Tissue staining was performed on the slide.

All washes were performed with TBS or 0.04% Triton X-100 in TBS (TBS-T 0.04%). Tissue was permeabilized with 0.4% Triton X-100 in TBS (TBS-T) for 30 minutes followed by 2 hours of 0.3M glycine in TBS-T with 10% normal donkey serum (NDS) and 3% bovine serum albumin (BSA) to quench free aldehydes. Primary incubation with antibodies diluted in TBS-T + 10% NDS + 3% BSA was done for 24-48 hours in 4C. Secondary antibodies were diluted in TBS-T + 3% BSA and incubated for 2 hours at room temperature. Slices were then washed and incubated with DAPI or Hoescht stain for 5 minutes at room temperature before mounting on glass slides with ProLong Diamond Antifade Mountant (#P36970, Thermo Fisher Scientific) and a coverslip.

### Antibody Table

| Antibody | Catalog # | Concentration |
| --- | --- | --- |
| Cc3 | CST 9661 | 1:200 |
| EphA4 | AF641 | 1:50 |
| Foxp2 | AF5647 | 1:1000 |
| Foxp2 | AB1307 | 1:1000 |
| Foxp2 | AB16046 | 1:1000 |
| GFP | AB6556 | 1:500 |
| GFP | CS 2955 | 1:500 |
| GFP | AL GFP-110 | 1:500 |
| Ki67 | Inv14-5698-82 | 1:500 |
| NeuN | MAB337 | 1:200 |
| TdTomato | 16D7 EST203 | 1:1000 |

### Fluorescent In-situ Hybridization (smFISH)

Adult mice were sacrificed by cervical dislocation and rapid decapitation followed by brain removal and placed in ice-cold PBS before freezing rapidly with dry ice and stored at -80C. Brains were then mounted in OCT, frozen, and sliced to∼15um on slide. Slides were kept at -20C for ∼30 minutes before stored in -80C again. We used RNAscope from ACDBio to perform smFISH. We followed the standard fresh-frozen tissue processing protocol for RNAscope Fluorescent Multiplex assay. Slides were mounted and coversliped and stored in 4C until imaging.

| Probe | Catalog # |
| --- | --- |
| ms Foxp2 | 428791 |
| ms Cre | 312281 |
| ms Sst | 404631 |
| ms Slc17a6 | 319171 |
| ms Slc32a1 | 319191 |
| ms Ptprk | 504561 |

### Imaging and Image Analysis

Images were acquired using a Zeiss 710 or 880 confocal microscope at the UT Southwestern Neuroscience Microscopy Facility. Images were collected at 10x or 20x as Z-stacks and tile scans with 10% overlap. Raw images were processed by stitching tiles and collapsed with maximum intensity projection with Zeiss ZEN Lite before analysis with FIJI. For IHC image analysis, three non-overlapping representative 233 µm squares from each IC slide were chosen and manually counted for positive antibody signal and overlap per DAPI/Hoescht identified nuclei. In control ICs, a representative square was placed in the DCIC, LCIC, and CIC. Given the substantial reduction in IC size in *Foxp2* cKO mice, regional subdivision of the IC was not used for quantitative analysis of locating specific cell types. Instead, representative mid, dorsal, and lateral-IC coronal sections were selected, and cells positive for individual probes or probe combinations were quantified. For ISH image analysis, the IC was selected and nuclear regions, regions of interest (ROIs), selected by thresholding and watershedding. Each probe channel was thresholded and calculated for area percent positivity in each ROI. ROI was considered positive for a signal probe if the minimum fraction of ROI pixels positive was 1-5% depending on probe average coverage.

### Behavior

All behavior was performed on a mix of male and female P50-60 mice. The first day each mouse is weighed and underwent an open field assay. The second day consisted of the startle assay, followed by the PPI assay on the third day. Mice were then either underwent ABR, perfused for IHC, or brain dissected and flash frozen for smFISH.

#### Open Field

Adult mice of both genotypes and sex were placed into a square open field and motion tracked with EthoVision software for 10 minutes. The arena was segmented into periphery (10cm border along wall) and non-periphery. Average velocity, time spent in periphery, and total distance traveled was calculated.

#### Startle and Paired Pulse Inhibition

Adult mice of both genotypes and sex were assayed for auditory startle response and paired pulse inhibition to auditory startle. Mice are individually placed into a small plastic tube located in a sound-attenuating chamber. After 5 min habituation to the chamber and background noise (65-70 dB), startle eliciting auditory stimuli, or no stimuli, is played through speakers located in the chamber. The startle stimulus can vary from 80 to 120 dB (40 ms duration, white noise) with an average interstimulus interval of 20 sec (range 13 - 27 sec). For PPI testing, the animals are placed in the chambers as above. Mice are exposed to 10 startle only trials played as the first and last 10 trials. During the prepulse trials, a second tone (2 - 16 dB) above the background noise is played immediately preceding (100 msec) the 120 dB startle eliciting stimulus along with no stimuli or only startle stimuli trials. Prepulse sounds are at a final 72, 74, 78, 82 dBs, beneath the startle threshold (as measured during the startle trial). San Diego Instruments, SR-Lab Startle Response System. Startle and PPI data analysis was performed in R. Startle trials were averaged per mouse per trial condition. For PPI analysis, the first and last 10 startle only trials are ignored. Baseline per mouse movement average was calculated with the 40 ms no stimulus trials. PPI was calculated using the standard percent inhibition metric for each subject and prepulse condition. Statistical outliers were removed if a mouse was statistically identified as an outlier in 2 or more trial averages using IQR-based detection. Average time between stimuli and max velocity was also calculated similarly in both startle and PPI analysis. A repeated-measures linear mixed-effects model was performed with genotype and prepulse condition as the between subject factor and within-subject factor, respectively. Subject was included as a random intercept to account for repeated measurements. Estimated marginal means computed from the fitted model was used to find genotype differences within each prepulse condition with emmeans post-hoc contrasts.

#### Auditory Brainstem Response

Adult mice were anesthetized with (dosage) ketamine/xylazine, placed on a warming pad and electrodes were placed subcutaneously below the ear for reference, midline on the head for the channel and below the contralateral ear for ground. Testing was performed in a sound attenuating chamber (Whisper Room Inc.) with equipment from Tucker Davis Technologies: RZ6-A-P1 processor, Medusa4Z amplifier, MF1-M speakers, and DPM microphone system. Using BioSigRZ software, ABR waveforms (averaged from 512 repetitions) were acquired for clicks (0.1msec, rate 21/sec) and for frequency specific (4, 8,16,24,32 kHz) tone bursts (5msec, rate 21/sec) at sound pressure levels of 20-90dB (10dB increments). Wave amplitudes were measured by the voltage difference between the positive (P) and negative (N) peaks for each wave at the 90dB click stimulus level.

### Electrophysiology

#### Electroencephalogram (EEG) recordings

For EEG recordings, we performed electrocorticography (ECoG) as described previously ^81,82^. Recordings were made from screw electrodes which were implanted into the skull, touching the dural surface over frontal and auditory regions of the cortex (see location details below). Mice were habituated for 15–20 min after being brought into the EEG recording room. The mice were then placed in an anechoic foam-lined soundproof chamber and tethers were connected to the implanted posts through a 3-channel tether that was passed through a commutator located directly above the recording cage. Mice were habituated to being tethered to the commutator for an additional 15–20 min. The tethers were connected to a 16-channel S-Box channel splitter which fed data through a PZ3 low impedance amplifier (Tucker-Davis Technologies, TDT, Alachua, FL). The EEG data is recorded through a RZ2 Bio Amp Processor (TDT, Alachua, FL). After habituation, at least 5 min of resting EEG were recorded in the absence of auditory stimuli. Next, the mice were presented with 300 presentations of a chirp-modulated tone (1–100 Hz, 2 s). After recordings were completed, mice were returned to the colony. The lead to the occipital cortex served as a reference electrode for the frontal and auditory cortex screws. Acquisition hardware was set to high-pass (>0.5 Hz) and low-pass (<150 Hz) filters. Data were sampled at a rate of 610.35 Hz via OpenProject software (TDT, Alachua, FL). Movement is known to elevate cortical gain during EEG recordings ^83^. To account for movement, we placed a piezoelectric transducer below recording cages to track movement during recordings. Prior to collecting experimental data, the activity from the piezoelectric transducer was evaluated in conjunction with video (RV2 Video Capture System, TDT, Alachua, FL) to confirm threshold criteria for movement. The EEG segments in which movement was occurring were rejected and excluded from analysis for resting EEG recordings and non-phase locked power measurements.

#### Surgery for EEG recordings

Mice were anesthetized using 1–2% isoflurane and this was maintained through a nose-mask attached to a stereotaxic frame (Narishigi Group, Japan). Artificial tears were used to keep the eyes moist during anesthesia and doses of buprenorphine sustained release (SR) and carprofen were administered subcutaneously. Toe/tail pinch reflexes were monitored throughout. Once anesthetized, a midline incision was made in the scalp to expose the skull. A precision micro-drill (Fine Science Tools, Foster City, CA) was used to create three 0.9 mm holes over the right frontal cortex (+3.0 mm, +1.6 mm | Coordinates relative to bregma: anterior/posterior, medial/lateral), right auditory cortex (−1.6 mm, +4.8 mm), and the left occipital (−4.2 mm, −5.1 mm). A 3-channel post (MS333–2-A-SPC, P1 Technologies, Roanoke VA) was connected to screws (8L003905201F, P1 Technologies) which were placed into the drilled holes until they contacted the dura mater. Dental cement (Panavia SA cement plus, Kuraray America, Houston, Texas) was applied to cover the screws, base of the 3-channel post, and skull. Hardening of the cement was expedited using a LED curing light. Mice were then placed on a circulating water heating pad and given 0.5 mL saline to aid in recovery. After recovery from surgery, the mice were individually housed and were returned to the vivarium and monitored daily until the day of recordings. Additional doses of carprofen were given 24 h and 48 h post-surgery. Mice were given between three to five days of post-operative recovery time and recordings were conducted between P50-P60.

#### Auditory stimulus presentation for EEG

Auditory stimuli were generated using Real-Time Processor Visual Design Studio (RPvdsEx) software and delivered using a RZ6 Multi I/O Processor (Tucker Davis Technologies, Alachua, FL). For the presentation of auditory stimuli, a free-field speaker (MF-1 speakers, TDT, Alachua, FL) was mounted ∼12 in. directly above a cage with fresh bedding. This was housed inside a chamber (Expanded PVC S.A.C. w/window, Med Associates, Inc.) surrounded by a custom-made Faraday cage. Both sound level measurements and recordings to confirm intended generation of waveforms were performed by the same system (CM16/CMPA microphones, 416H 200 amplifier, Avisoft-Bioacoustics) by placing the microphone on the floor of the recording chamber. All sounds were created and presented at a sampling rate of 97,948 samples/s. We recorded event related potentials (ERP). Pulse train stimuli consisted of a sound pulse train of four broadband white noise “clicks” (50 ms duration, separated by 500 ms with a 4 s interval between each train, 70 ± 3 dB SPL). Responses were collected from 100 trials. Briefly, a “chirp” is a stimulus in which the carrier sound amplitude is modulated by a sinusoid whose frequency increases from 1 to 100 Hz. This is preceded by a linear volume ramp to avoid onset responses contaminating phase locking to the amplitude modulation of the chirp. The chirp was repeated for 300 trials. Including the prestimulus 1 s and post-stimulus 0.5 s baseline periods for each recorded trial, each trial was 4.5 s long, and the inter-trial interval was 3 s. Previous studies revealed similar phenotypes when the modulation was in an upward (1 to 100 Hz) or downward (100 to 1 Hz) direction ^82^. Therefore, just the upward modulation was studied here.

#### Electrophysiology data analysis

EEG data were analyzed in custom LABVIEW software (National Instruments Corp, Austin, TX). Data were pre-processed using a 0.5-150 Hz bandpass filter. A 60 Hz notch filter was applied. All recordings were cleaned for artifacts using a semi-automatic procedure in LABVIEW. A similar semi-automatic procedure was used to define movement based on signals from piezoelectric transducers. ITPC and raw power as a function of time were calculated as described previously ^84^. For each mouse, traces from individual trials involving sound stimuli (e.g. chirp) underwent a Morelet wavelet transformation, *F_k_(f,t)*, which refers to the complex wavelet coefficient at a given frequency and time for the kth trial. The PLF as a function of time and frequency was derived from calculating intertrial phase coherence (ITPC). The Morelet wavelet transforms from all trials were used to calculate the ITPC as follows:

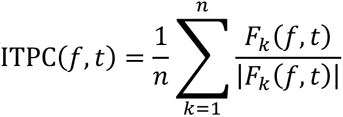

where f is the frequency, t is the time point, and n is total trial number. We used a peak frequency to width ratio of 15. Raw power as a function of time and frequency was calculated by obtaining a time-series power spectrum. This was done by extracting absolute values from complex values obtained from the Morelet wavelet transformation (squareroot[real^2^ + imaginary^2^]) for each cell in each trial matrix. Absolute value matrices were then averaged for all trials for a given mouse, and group grand average matrices were then compiled for each group. To perform statistical comparisons of these data, we performed a nonparametric cluster analysis in MATLAB R2020a (MathWorks, Natick, MA). These data have 2 dependent variables, frequency and time, and consequently, we obtain a 2 dimensional matrix of values from each mouse. The cluster analysis was used to determine contiguous regions in the matrix that were significantly different from a distribution of 1000 randomized Monte Carlo permutations based on previously published methods 25. In this analysis, if cluster sizes of the real genotype assignments (both positive and negative direction, resulting in a two-tailed alpha of p = 0.025) were larger than 97.25% of the random group assignments, those clusters were considered significantly different between genotypes. This method avoids statistical assumptions about the data and avoids the multiple comparison problem. We have used this analysis for the exact same type of EEG data obtained from mice in previous publications ^85,86^. Differences in EEG waveforms in the time domain were calculated by detecting time epochs where average plus/minus 95% confidence limits were nonoverlapping. p values < 0.05 were considered significant for ANOVA. In all cases where genotype means are reported, SEM was used. Statistical analyses were performed using the program GraphPad Prism 9.5.1. All analysis was performed blinded to genotype.

#### Brain slices and Recordings

We used mice aged postnatal day (P) 14-20 for the electrophysiological measurements (No difference in average age between genotypic groups). We used younger tissue for its increased viability and to avoid obstruction of dense extracellular matrix that exists in adult. Mice were anesthetized with a combination of ketamine (125 mg/kg) and xylazine (25 mg/kg), and brains were quickly dissected and transferred to partially frozen dissection buffer ^87^. 300 μm thick brain slices containing the dorsal striatum were prepared using a Vibratome (Leica VTS 1200S) and immediately placed in nominal artificial cerebrospinal fluid (nACSF) (bubbled with a mixture of 5% CO2 and 95% O2), first at 35°C for 30 min, and then at room temperature for at least 30 min before commencement of recordings ^87^. We used glass pipettes of 5-8 MΩ resistance to perform whole cell recordings of Cre-expressing neurons, which were identified based on the expression of a YFP reporter. We included neurons with a maximum acceptable resting membrane potential of -50 mV, capacitance greater than 15 pF, stable baseline current within 15 pA, and a series resistance of less than 25 MΩ in our analysis. The average series resistance did not exhibit significant differences between control and *Foxp2* cKO neurons. The junction potential was ∼10 mV and was not corrected. All recordings were performed at a temperature of 26 °C. Unless stated otherwise, we recorded and analyzed all electrophysiological parameters using custom software (Labview; National Instruments, Austin, TX; https://github.com/ColdP1228/Custom-Data-Acquisition-Program-20; copy archived at https://github.com/elifesciences-publications/Custom-Data-Acquisition-Program-).

Input resistance was measured at a membrane potential of −70 mV. For sEPSC recordings, neurons were voltage clamped at −70 mV and spontaneously occurring synaptic events were recorded in 10 traces, each lasting 10 s. Data analysis was performed using an automatic detection program (MiniAnalysis; Synaptosoft Inc, Decatur, Ga.) with a detection threshold at a value exceedingly at least 5 standard deviations (10 pA) of the baseline noise. Additionally, we visually confirmed the detected events. The detection threshold remained constant throughout each recording.

#### Brain slice Electrophysiology Solutions

We prepared 250µm brain slices in a semi-frozen dissection buffer (bubbled with 95% O2 and 5% CO2) containing the following components (in mM): 110 Choline-Cl, 25 Dextrose, 25 NaHCO3, 3 Ascorbic Acid, 3.1 Sodium Pyruvate, 2.5 KCl, 1.25 NaH2PO4, 7 MgCl2, 0.5 CaCl2.2H2O, and 0.2 Kyneuric Acid, with a pH of 7.3-7.4 and an osmolality of 280-285 mOsm. The slices were transferred to nACSF that contained (in mM): 125 NaCl, 25 NaHCO3, 10 Dextrose, 3 KCl, 1.25 NaH2PO4, 2 MgCl2, and 2 CaCl2.2H2O with a pH of 7.3-7.4 and an osmolality of 300-305 mOsm, saturated with 95% O2/5% CO2. The pipette (internal) solution consisted of (in mM): 130 K-Gluconate, 10 HEPES, 6 KCl, 3 NaCl, 0.2 EGTA, 14 phosphocreatine-Tris, 4 ATP-Mg and, 0.4 GTP-Tris with a pH of 7.25 and an osmolality of 300 mOsm.

### Paired Single-Nucleus RNA and ATAC-seq

#### Tissue Processing and Library Generation

Protocol is derived from prior lab publications ^88^ and the 10X Genomics Multiome tissue nuclei isolation protocol (10X Genomics CG000375). Mice who provided tissue were sacrificed by cervical dislocation and rapid decapitation. Brains were quickly removed and placed in ice-cold PBS, sliced into 1 mm sections on ice-cold metal brain matrix, and had their IC dissected from relevant slices. Tissue was flash frozen in liquid nitrogen for storage. Final data consists of 3 samples per genotype per timepoint with both males and females. Each processed sample consisted of 2 pooled animals of identical sex and genotype. Nuclei isolation began with tissue disassociation suspended in ice-cold lysis buffer with 2 mL Dounce homogenizers. Pestle A was used for 15-20 times followed by 15-20 times with pestle B. Sample was then transferred to ice-cold homogenization buffer. After centrifuging the sample, supernatant was removed and replaced with nuclei suspension buffer (NSB). We performed a sucrose gradient spun in a centrifuge for 60 minutes at 4C. Supernatant was removed and replaced with new NSB to be filtered through 50 µm Flowmi Cell strainer. An aliquot of sample nuclei was mixed with trypan blue at 1:1 and counted on DHC-F01 counting chip. If quality and quantity of nuclei is sufficient, sample was centrifuged, supernatant removed and sample resuspended in diluted nuclei buffer. Sample libraries were created via 10X Genomics Multiome kit and protocol. Quality of sample was checked with Aligent TapeStation. Quantity of final sample was measured with Qbit before pooling samples and sequencing. Libraries were sequenced by the McDermott Sequencing Core at UTSW with Illumina NextSeq 2000s sequencer. Unprocessed sequencing data were provided as binary base call (BCL) files. BCL files were de-multiplexed with the 10X Genomics index using Illumina’s bcl2fastq and mkfastq command from 10x Genomics Cell Ranger-ARC v2.0.2 tools. Extracted paired-end fastq files were check for read quality using FastQC v0.11.5. CellRanger-ARC v2.0.2 count was used to align reference genome GRCm39 to sample libraries.

#### Quality Control

The raw gene-count matrix containing cells as rows and genes as columns was used for ambient RNA removal with CellBender v0.3.0 remove-background. DoubletFinder v2.0.4 was used to predict and remove doublets. Data was filtered by removing clusters with an average intronic read ratio below 50% and a mitochondrial DNA percentage above 10%. Clusters dominated by single samples were also removed. Genes from mitochondrial chromosome and chromosomes X and Y were removed. In the adult dataset, clusters identified as microglia, oligodendrocyte precursor cells, and immature oligodendrocytes were removed due to low nuclei number. Next, we wanted to exclude non IC nuclei from the dataset that may have been captured during dissection. To identify what cells could have come from adjacent superior colliculus (SC) or periaqueductal grey (PAG) regions, we used publicly available data to identify clusters contributed to by surrounding regions. Recent databases have been released of mouse whole brain cellular atlases, combining single cell RNA-seq and spatial transcriptomics to identify cell types by region and gene expression ^89^. Using the MapMyCells online tool (https://portal.brain-map.org/atlases-and-data/bkp/mapmycells) ^90^, we were able to identify neuronal clusters with significant (>90%) proportion of SC or PAG neurons and remove them from the analysis. One cluster showed a mixed population, with only 12% of cells mapped to IC glutamatergic neurons. These neurons were kept and the rest discarded, as they represented a potential subcluster of IC neurons. The final remaining nuclei were then normalized and reclustered before annotation.

### Data Analysis

#### Clustering analysis

The snRNA-seq dataset was analyzed using the Seurat/Signal analysis pipeline in R. Data were log-normalized with NormalizeData and regressed for covariates as described in the Seurat analysis pipeline. Data were used to calculate principal components (PCs) and identify top variable genes. With an elbow plot and jackstraw analysis, statistically significant PCs were used to identify clusters within the data using the original Louvain algorithm. Seurat RPCA integration was used to regress for batch effects.

#### Cell type annotation

Cluster marker genes were identified using FindAllMarkers with receiver operating characteristic (ROC) test. Our cluster markers were compared with general neuronal and glial cell markers. For control cell type excitatory (*Vglut2*+) and inhibitory (*Gad1*+ and *Gad2*+) clusters were subsetted and reclustered. Gene expression matricies were exported for Allen Brain Map MapMyCells online tool. Input data was compared to the Allen Institute Mouse Whole Brain dataset using hierarchical correlation mapping. Clusters with >90% of nuclei mapping to IC glutamatergic (181 IC Tfap2d Maf Glut) or GABAergic (198 IC Six3 En2 Gaba) were kept and reclustered. Cluster resolution was chosen based on cluster separation and representation by control and *Foxp2* cKO datasets. Marker genes were filtered for high significance and transcription factor identity. LabelTransfer was used to check *Foxp2* cKO transcriptomic similarity to control cells in juvenile clusters before transferring annotations. R package clustree was used to determine relationships between clusters at variable resolutions. We transferred high-confidence control adult neuronal annotations to the juvenile dataset using LabelTransfer on normalized, clustered data. Non-IC clusters of nuclei were also identified and removed in the same manner as the adult dataset.

#### Differential abundance analysis

To analyze cell proportions between genotypes within clusters, we used scCoda v0.1.9. Clusters were considered significantly differentially abundant between genotypes using sample counts if FDR < 0.05. Milo differential analysis was also used on the adult dataset using a KNN algorithm. Clusters were considered significantly differentially abundant between genotypes if the spatial FDR < 0.05.

#### Peak Mapping

The paired snATAC-seq dataset retained the snRNA-seq derived annotations. Peaks were called using Signac CallPeaks function, which wraps the MACS2 function to call peaks.

#### Motifs

To determine overrepresented motifs in a set of significant differentially accessible peaks, DARs, we added motif information using AddMotifs function from mm38 in JASPAR2020 database. Then with FindMotifs, we computed motif enrichment compared to the background using the function’s hypergeometric test with Benjamini-Hochberg p adjust method. Significant enrichment was defined as FDR < 0.05 and fold enrichment > log_2_(1.2).

#### Differentially expressed gene (DEG) and differentially accessible region (DAR) analyses

To determine differentially expressed genes, cell types were pseudobulked by cluster and genotype identity. Sample libraries with less than 5e^4^ features, clusters with less than 10 cells per sample, and groups with less than 2 samples per group were removed from analysis. Covariates included in analysis were sample, sex, and sequence batch. With edgeR, differential gene analysis on pseuodobulked clusters between genotypes was performed. Within each cluster, raw counts were aggregated at the sample level, enabling statistical inference across biological replicates rather than individual nuclei. Sample-cluster combinations with insufficient nuclei were excluded to ensure robust estimation. Differential gene expression between control and *Foxp2* cKO samples was then assessed within each cluster using edgeR, with genotype modeled as a fixed effect and library size normalization, dispersion estimation, and sex and sequencing batch covariates included in the analysis. Differentially expressed genes were identified based on false discovery rate-corrected significance (FDR =< 0.05) and fold-change thresholds (log_2_FC >= 1.2).

#### Gene Ontology

Gene ontology analysis was performed in R using enrichR with databases GO Biological Processes, Molecular Function, and Cellular Compartment 2025, and SynGO 2024. Terms were considered significant if they had a corrected p-value <0.05 and a minimum of three genes. High-confidence ASD associated genes were downloaded from the SFARI-ASD database. NDD associated genes in the Definitive category were downloaded from the SysNDD database. Significant overlap was determined using the hypergeometric overlap test.

## Author contributions

Study design: MJ, JRG, GK

Investigation: MJ, YTH, AH, SK, AK, HS

Formal analysis: MJ

Supervision: JRG, GK

Writing—original draft: MJ

Writing—review and editing: MJ, JRG, GK

## Supplementary Materials

Figures S1 to S5

Table S1

## Acknowledgements

We thank the UTSW Neuroscience Microscopy Facility for assistance with imaging and the UTSW Rodent Behavior Core for assistance with behavioral assays.

## AI Use Disclosures

Text was edited for grammar and clarity, and code used for analysis was in part generated and edited using publicly available artificial intelligence language learning models.

---

**Supplementary Figure 1:**
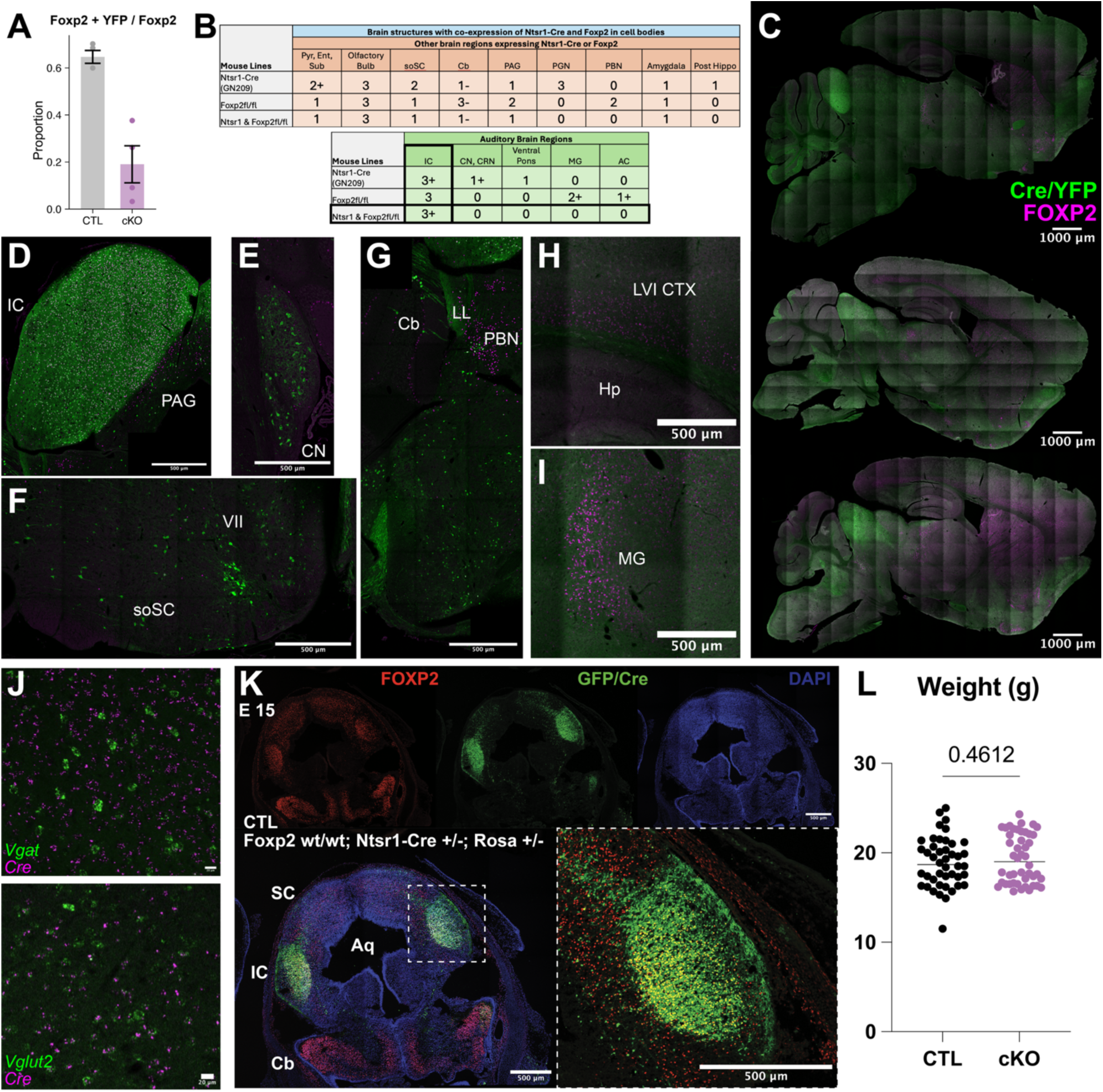
Overlapping expression of *Foxp2* and Ntsr1-Cre GN209. Within the auditory system, including the cortex and thalamus, the major overlap of Cre and *Foxp2* expression occurs in the IC. (A) Percent of FOXP2+YFP co-expressing cells in the FOXP2 population in the IC. (B) Table of Cre and *Foxp2* expression. 0-3 expression strength, +/- designates excitatory/inhibitory neurons. (C) Sagittal sections of control brain of Cre/YFP and FOXP2 expression. (D) Inferior colliculus (IC) and peri-aqueductal gray (PAG). (E) Cochlear nucleus (CN). (F) Superior olivary complex (soSC). (G) Cerebellum (Cb), lateral lemniscus (LL), and peribrachial nucleus (PBN). (G) Layer 6 of cortex (LVI CTX) and hippocampus (Hp). (I) Medial geniculate body (MG). (J) ISH of control IC central nucleus with *Vglut* (top) and *Vglut2* (bottom) co-labeled with *Cre*. (K) IHC of coronal E15 control slice showing expression and overlap of FOXP2 (red) and YFP/Cre (green) in the developing dorsal tectum. (L) Average weights of adult control and *Foxp2* cKO mice.

**Supplementary Figure 2:**
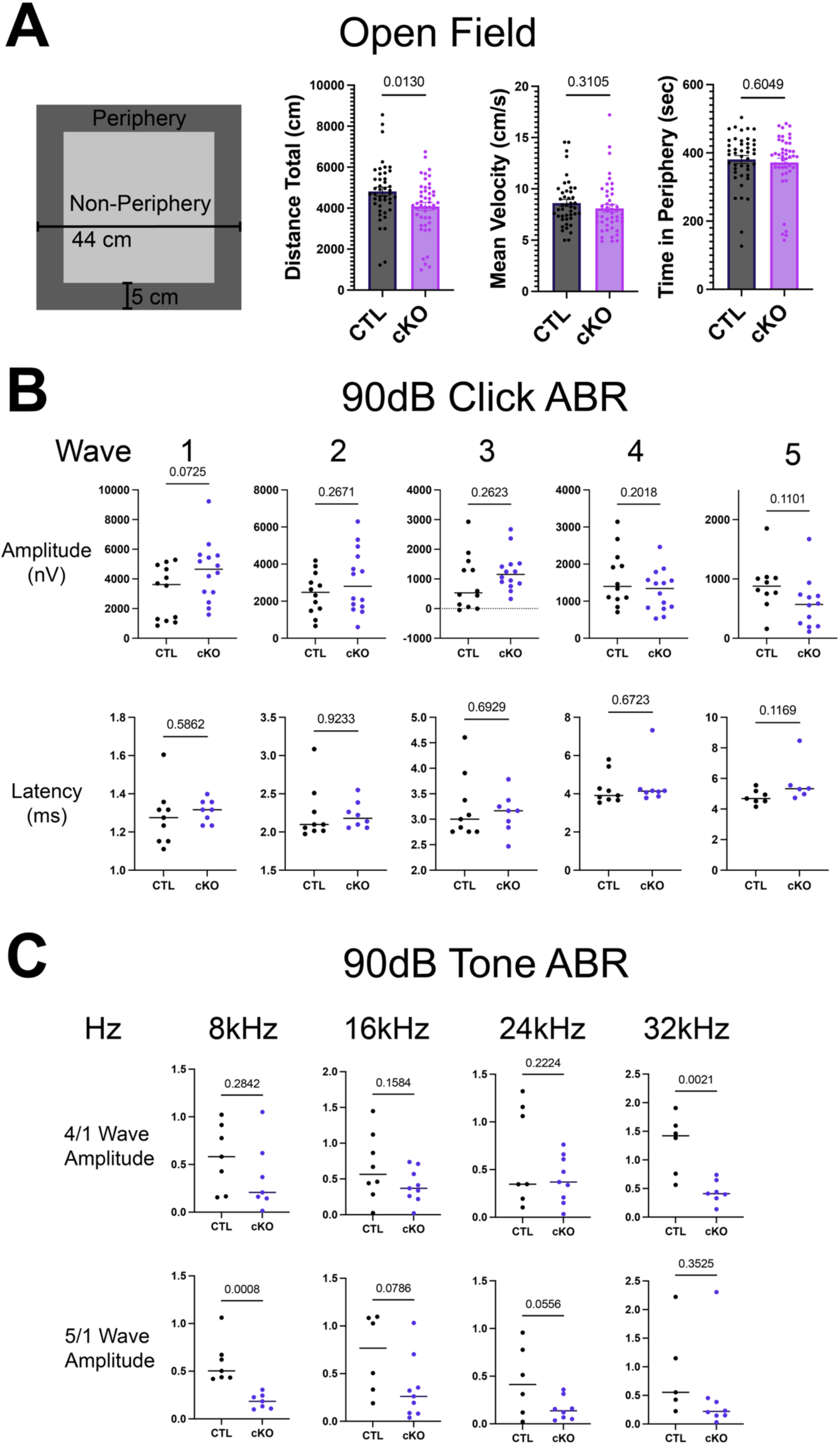
Additional behavior and ABR measurements. (A) Adult control and *Foxp2* cKO mice were placed in an open field and motion tracked for 10 minutes. Diagram of arena (left), and total distance, mean velocity, and time in periphery (right) were measured. (B) Average amplitude (top row) and latency (bottom row) to each ABR wave per mouse in click trials. (C) 4/1 (top row) and 5/1 (bottom row) wave amplitude averages per mouse during tone trials.

**Supplementary Figure 3:**
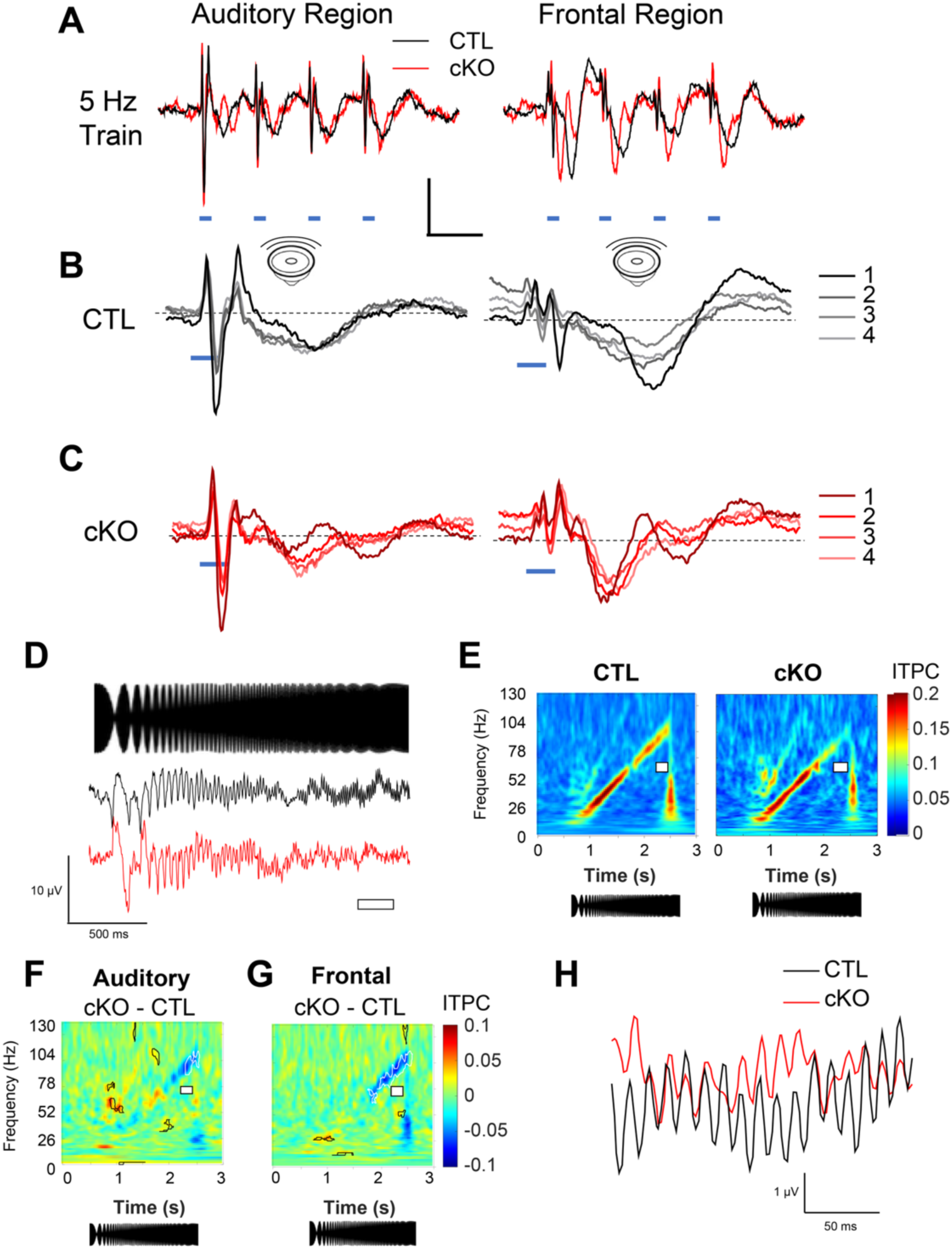
EEG results. A-C) 4 average responses to the 4 click sounds are plotted for the control and *Foxp2*. D-H) Auditory cortex activity in response to a chirp sound is more weakly time-locked to higher frequencies in the *Foxp2* IC-cKO. A) Average activity measured during the 4 pulse train for control and *Foxp2* cKO mice in auditory and frontal cortical regions. B,C) all 4 average pulses for each sound in the train. D) The chirp sound was a 14 kHz sinewave modulated in amplitude starting at 1 Hz and gradually increasing to 100 Hz (top). Average traces over all controls and cKO mice (below). E) Average plots for ITPC time-course at the auditory electrode. Plot includes the last 0.5 seconds of the initial ramp. F,G) Average difference plots for auditory (F) and frontal electrodes (G). H) Average ECoG signal for control and *Foxp2* cKO during an interval where control has statistically larger ITPC (interval marked by the white box in E, F, and G). Bold lines in (F) and (G) mark statistically different regions of the plots (p<0.05). Black and white lines represent increases and decreases, respectively.

**Supplementary Figure 4:**
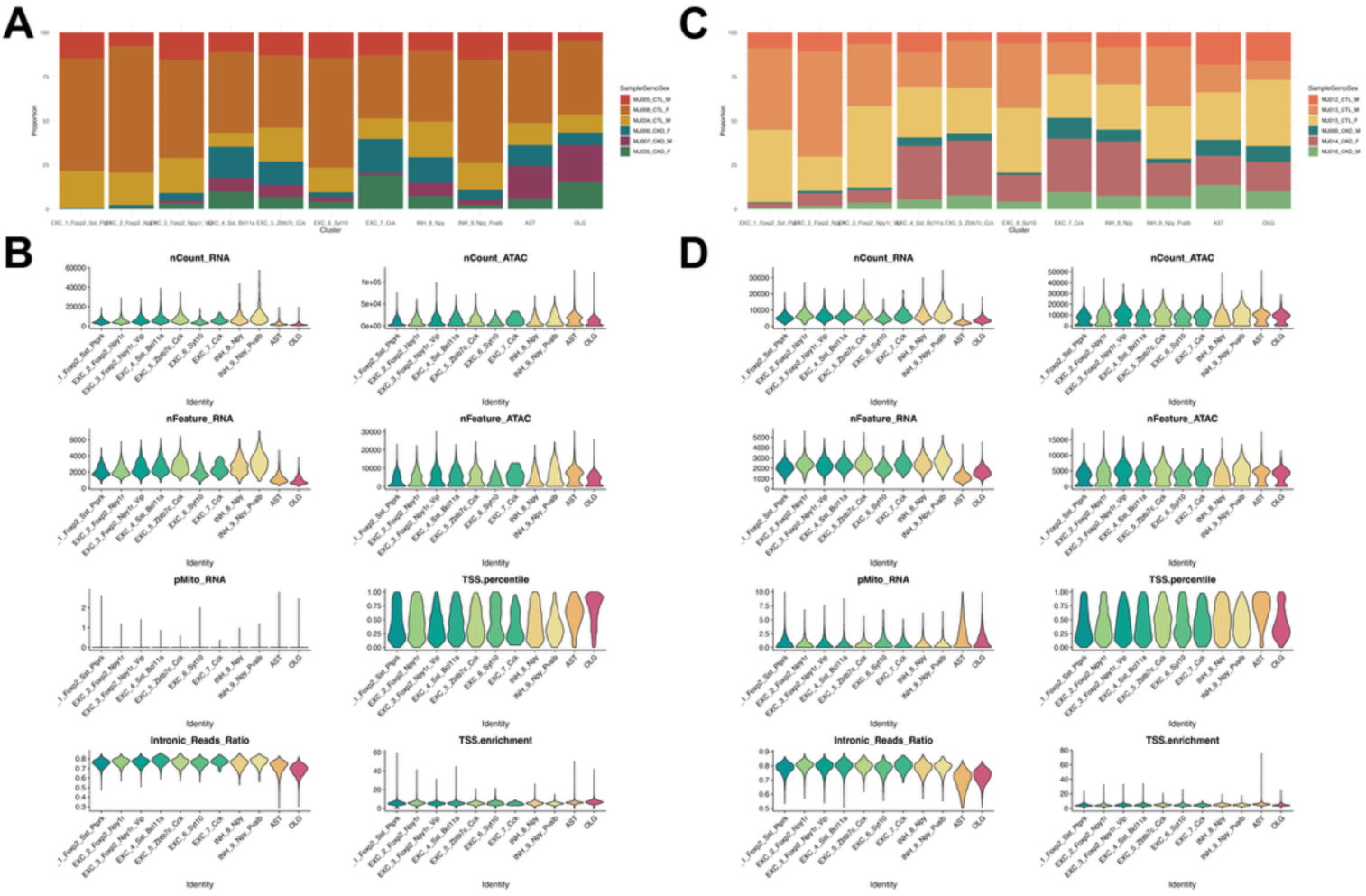
Multiome QC. A) Stacked bar plots showing the composition of each cell type by sample in the adult dataset. (B) Quality control metadata values per cell type in the adult dataset. (C, D) Same for juvenile dataset.

**Supplementary Figure 5:**
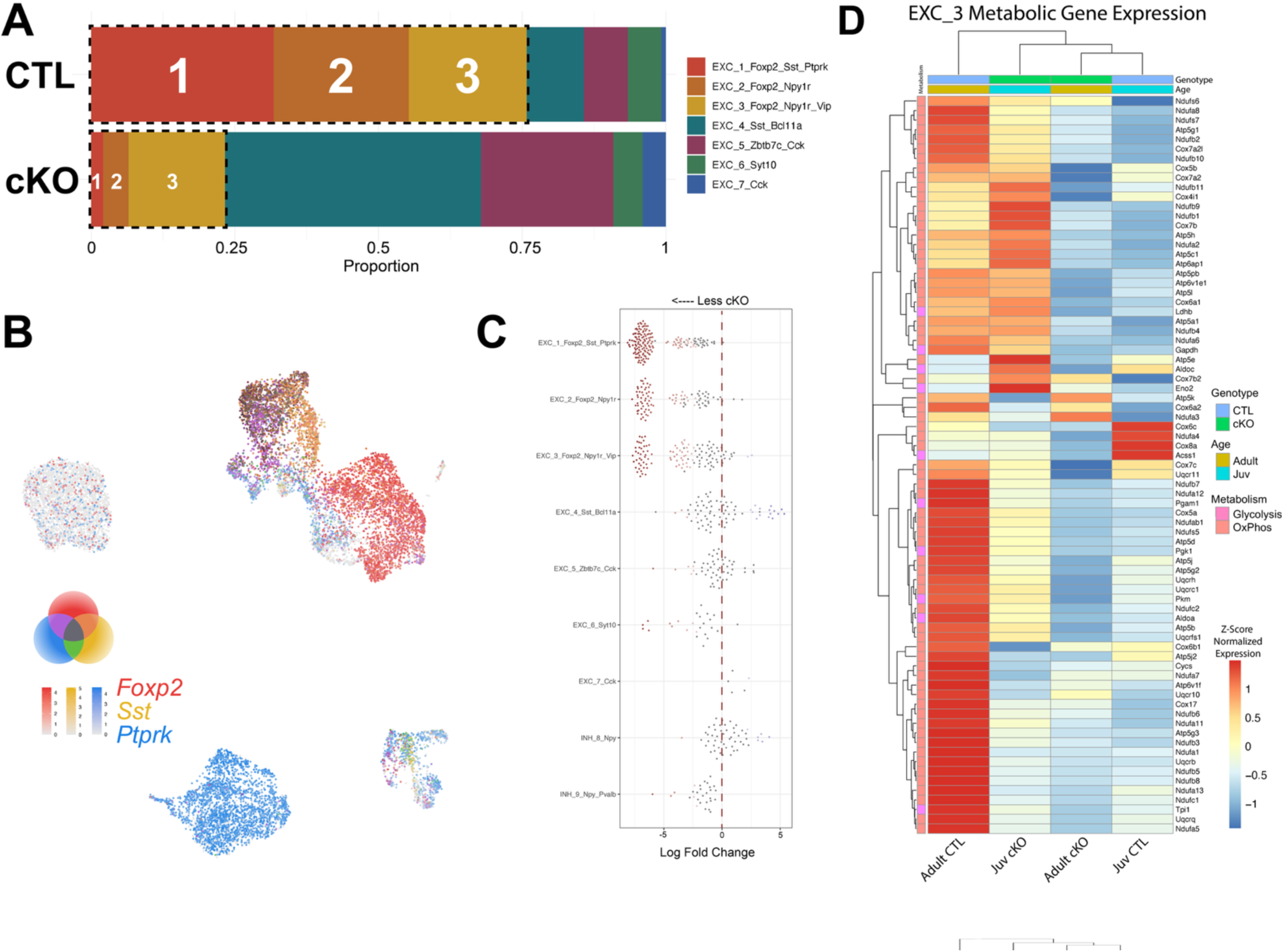
Adult IC neural subtypes and metabolism. A) Bar graph of excitatory neuron subtype proportions in control and *Foxp2* cKO adult data. High *Foxp2* expressing clusters are outlined in black dashed lines. B) Feature plot of the adult control UMAP showing normalized expression of *Foxp2* (red), *Sst* (yellow), and *Ptprk* (blue). C) Proportion of neuronal cell types between genotypes with Milo differential abundance analysis, with significance indicated in red, spatial FDR < 0.05. D) Z-score normalized expression of metabolism genes between adult and juvenile control and *Foxp2* cKO integrated data in cluster EXC 3.

## Supplementary Tables

**Supplementary Table 1: Multiome QC meta, Adult and Juvenile DEGs, DARs, Motifs**

## References

1. Batra, R. (2014). Inferior Colliculus. In Encyclopedia of the Neurological Sciences (Elsevier), pp. 692–694. 10.1016/B978-0-12-385157-4.01153-2.

2. Winer, J.A., and Schreiner, C. eds. (2005). The inferior colliculus (Springer).

3. Auditory System (2015). In The Rat Nervous System (Elsevier), pp. 865–946. 10.1016/b978-0-12-374245-2.00029-2.

4. Ito, T., Ono, M., and Oliver, D.L. (2018). Neuron Types, Intrinsic Circuits, and Plasticity in the Inferior Colliculus. In The Oxford Handbook of the Auditory Brainstem, K. Kandler, ed. (Oxford University Press), pp. 503–526. 10.1093/oxfordhb/9780190849061.013.25.

5. Lai, C.S.L. (2003). FOXP2 expression during brain development coincides with adult sites of pathology in a severe speech and language disorder. Brain 126, 2455–2462. 10.1093/brain/awg247.

6. Gurung, B., and Fritzsch, B. (2014). Time Course of Embryonic Midbrain and Thalamic Auditory Connection Development in Mice as Revealed by Carbocyanine Dye Tracing.

7. Liu, M., Wang, Y., Jiang, L., Zhang, X., Wang, C., and Zhang, T. (2024). Research progress of the inferior colliculus: from Neuron, neural circuit to auditory disease. Brain Res. 1828, 148775. 10.1016/j.brainres.2024.148775.

8. Goyer, D., Silveira, M.A., George, A.P., Beebe, N.L., Edelbrock, R.M., Malinski, P.T., Schofield, B.R., and Roberts, M.T. A novel class of inferior colliculus principal neurons labeled in vasoactive intestinal peptide-Cre mice.

9. Peruzzi, D., Sivaramakrishnan, S., and Oliver, D.L. (2000). Identification of cell types in brain slices of the inferior colliculus. Neuroscience 101, 403–416. 10.1016/S0306-4522(00)00382-1.

10. Shu, W., Yang, H., Zhang, L., Lu, M.M., and Morrisey, E.E. (2001). Characterization of a New Subfamily of Winged-helix/Forkhead (Fox) Genes That Are Expressed in the Lung and Act as Transcriptional Repressors. J. Biol. Chem. 276, 27488–27497. 10.1074/jbc.M100636200.

11. Kast, R.J., Lanjewar, A.L., Smith, C.D., and Levitt, P. (2019). FOXP2 exhibits projection neuron class specific expression, but is not required for multiple aspects of cortical histogenesis. eLife 8, e42012. 10.7554/eLife.42012.

12. ADHD Working Group of the Psychiatric Genomics Consortium (PGC), Early Lifecourse & Genetic Epidemiology (EAGLE) Consortium, 23andMe Research Team, Demontis, D., Walters, R.K., Martin, J., Mattheisen, M., Als, T.D., Agerbo, E., Baldursson, G., et al. (2019). Discovery of the first genome-wide significant risk loci for attention deficit/hyperactivity disorder. Nat. Genet. 51, 63–75. 10.1038/s41588-018-0269-7.

13. Salmón-Gómez, G., Suárez-Pinilla, P., Setién-Suero, E., Martínez-Asensi, C., and Ayesa-Arriola, R. (2025). FoxP2 and Schizophrenia: a systematic review. J. Psychiatr. Res. 190, 205–215. 10.1016/j.jpsychires.2025.07.016.

14. Satterstrom, F.K., Kosmicki, J.A., Wang, J., Breen, M.S., De Rubeis, S., An, J.-Y., Peng, M., Collins, R., Grove, J., Klei, L., et al. (2020). Large-Scale Exome Sequencing Study Implicates Both Developmental and Functional Changes in the Neurobiology of Autism. Cell 180, 568–584.e23. 10.1016/j.cell.2019.12.036.

15. Lai, C.S.L., Fisher, S.E., Hurst, J.A., Vargha-Khadem, F., and Monaco, A.P. (2001). A forkhead-domain gene is mutated in a severe speech and language disorder. Nature 413, 519–523. 10.1038/35097076.

16. Zhang, S., Zhao, J., Guo, Z., Jones, J.A., Liu, P., and Liu, H. (2018). The Association Between Genetic Variation in FOXP2 and Sensorimotor Control of Speech Production. Front. Neurosci. 12, 666. 10.3389/fnins.2018.00666.

17. Vernes, S.C., Nicod, J., Elahi, F.M., Coventry, J.A., Kenny, N., Coupe, A.-M., Bird, L.E., Davies, K.E., and Fisher, S.E. (2006). Functional genetic analysis of mutations implicated in a human speech and language disorder. Hum. Mol. Genet. 15, 3154–3167. 10.1093/hmg/ddl392.

18. Mizutani, A., Matsuzaki, A., Momoi, M.Y., Fujita, E., Tanabe, Y., and Momoi, T. (2007). Intracellular distribution of a speech/language disorder associated FOXP2 mutant. Biochem. Biophys. Res. Commun. 353, 869–874. 10.1016/j.bbrc.2006.12.130.

19. Medvedeva, V.P., Rieger, M.A., Vieth, B., Mombereau, C., Ziegenhain, C., Ghosh, T., Cressant, A., Enard, W., Granon, S., Dougherty, J.D., et al. (2019). Altered social behavior in mice carrying a cortical Foxp2 deletion. Hum. Mol. Genet. 28, 701–717. 10.1093/hmg/ddy372.

20. Co, M., Hickey, S.L., Kulkarni, A., Harper, M., and Konopka, G. (2020). Cortical Foxp2 Supports Behavioral Flexibility and Developmental Dopamine D1 Receptor Expression. Cereb. Cortex N. Y. NY 30, 1855–1870. 10.1093/cercor/bhz209.

21. Druart, M., Groszer, M., and Le Magueresse, C. (2020). An Etiological Foxp2 Mutation Impairs Neuronal Gain in Layer VI Cortico-Thalamic Cells through Increased GABAB/GIRK Signaling. J. Neurosci. Off. J. Soc. Neurosci. 40, 8543–8555. 10.1523/JNEUROSCI.2615-19.2020.

22. Urbanus, B.H.A., Peter, S., Fisher, S.E., and De Zeeuw, C.I. (2020). Region-specific Foxp2 deletions in cortex, striatum or cerebellum cannot explain vocalization deficits observed in spontaneous global knockouts. Sci. Rep. 10, 21631. 10.1038/s41598-020-78531-8.

23. French, C.A., Vinueza Veloz, M.F., Zhou, K., Peter, S., Fisher, S.E., Costa, R.M., and De Zeeuw, C.I. (2019). Differential effects of Foxp2 disruption in distinct motor circuits. Mol. Psychiatry 24, 447–462. 10.1038/s41380-018-0199-x.

24. Groszer, M., Keays, D.A., Deacon, R.M.J., de Bono, J.P., Prasad-Mulcare, S., Gaub, S., Baum, M.G., French, C.A., Nicod, J., Coventry, J.A., et al. (2008). Impaired synaptic plasticity and motor learning in mice with a point mutation implicated in human speech deficits. Curr. Biol. CB 18, 354–362. 10.1016/j.cub.2008.01.060.

25. Enard, W., Gehre, S., Hammerschmidt, K., Hölter, S.M., Blass, T., Somel, M., Brückner, M.K., Schreiweis, C., Winter, C., Sohr, R., et al. (2009). A Humanized Version of Foxp2 Affects Cortico-Basal Ganglia Circuits in Mice. Cell 137, 961–971. 10.1016/j.cell.2009.03.041.

26. Schreiweis, C., Irinopoulou, T., Vieth, B., Laddada, L., Oury, F., Burguière, E., Enard, W., and Groszer, M. (2019). Mice carrying a humanized Foxp2 knock-in allele show region-specific shifts of striatal Foxp2 expression levels. Cortex 118, 212–222. 10.1016/j.cortex.2019.01.008.

27. Schreiweis, C., Bornschein, U., Burguière, E., Kerimoglu, C., Schreiter, S., Dannemann, M., Goyal, S., Rea, E., French, C.A., Puliyadi, R., et al. (2014). Humanized Foxp2 accelerates learning by enhancing transitions from declarative to procedural performance. Proc. Natl. Acad. Sci. 111, 14253–14258. 10.1073/pnas.1414542111.

28. Ahmed, N.I., Khandelwal, N., Anderson, A.G., Oh, E., Vollmer, R.M., Kulkarni, A., Gibson, J.R., and Konopka, G. (2024). Compensation between FOXP transcription factors maintains proper striatal function. Cell Rep. 43, 114257. 10.1016/j.celrep.2024.114257.

29. Chen, Y.-C., Kuo, H.-Y., Bornschein, U., Takahashi, H., Chen, S.-Y., Lu, K.-M., Yang, H.-Y., Chen, G.-M., Lin, J.-R., Lee, Y.-H., et al. (2016). Foxp2 controls synaptic wiring of corticostriatal circuits and vocal communication by opposing Mef2c. Nat. Neurosci. 19, 1513–1522. 10.1038/nn.4380.

30. Kuo, H.-Y., Chen, S.-Y., Huang, R.-C., Takahashi, H., Lee, Y.-H., Pang, H.-Y., Wu, C.-H., Graybiel, A.M., and Liu, F.-C. (2023). Speech- and language-linked *FOXP2* mutation targets protein motors in striatal neurons. Brain 146, 3542–3557. 10.1093/brain/awad090.

31. Usui, N., Co, M., Harper, M., Rieger, M.A., Dougherty, J.D., and Konopka, G. (2017). Sumoylation of FOXP2 Regulates Motor Function and Vocal Communication Through Purkinje Cell Development. Biol. Psychiatry 81, 220–230. 10.1016/j.biopsych.2016.02.008.

32. Takahashi, K., Liu, F., Oishi, T., Mori, T., Higo, N., Hayashi, M., Hirokawa, K., and Takahashi, H. (2008). Expression of *FOXP2* in the developing monkey forebrain: Comparison with the expression of the genes *FOXP1*, *PBX3*, and *MEIS2*. J. Comp. Neurol. 509, 180–189. 10.1002/cne.21740.

33. H. P. Brain Atlas, Midbrain, Foxp2 https://www.proteinatlas.org/ENSG00000128573-FOXP2/brain/midbrain#hpa_brain_rnaseqinferior_colliculus.

34. Campbell, P., Reep, R.L., Stoll, M.L., Ophir, A.G., and Phelps, S.M. (2009). Conservation and diversity of Foxp2 expression in muroid rodents: Functional implications. J. Comp. Neurol. 512, 84–100. 10.1002/cne.21881.

35. Kurt, S., Fisher, S.E., and Ehret, G. (2012). Foxp2 Mutations Impair Auditory-Motor Association Learning. PLoS ONE 7, e33130. 10.1371/journal.pone.0033130.

36. French, C.A., Groszer, M., Preece, C., Coupe, A.-M., Rajewsky, K., and Fisher, S.E. (2007). Generation of mice with a conditional Foxp2 null allele. Genes. N. Y. N 2000 *45*, 440–446. 10.1002/dvg.20305.

37. Kreeger, L.J., Connelly, C.J., Mehta, P., Zemelman, B.V., and Golding, N.L. (2021). Excitatory cholecystokinin neurons of the midbrain integrate diverse temporal responses and drive auditory thalamic subdomains. Proc. Natl. Acad. Sci. 118, e2007724118. 10.1073/pnas.2007724118.

38. Beebe, N.L., Silveira, M.A., Goyer, D., Noftz, W.A., Roberts, M.T., and Schofield, B.R. (2022). Neurotransmitter phenotype and axonal projection patterns of VIP-expressing neurons in the inferior colliculus. J. Chem. Neuroanat. 126, 102189. 10.1016/j.jchemneu.2022.102189.

39. Liu, M., Gao, Y., Xin, F., Hu, Y., Wang, T., Xie, F., Shao, C., Li, T., Wang, N., and Yuan, K. (2024). Parvalbumin and Somatostatin: Biomarkers for Two Parallel Tectothalamic Pathways in the Auditory Midbrain. J. Neurosci. 44, e1655232024. 10.1523/JNEUROSCI.1655-23.2024.

40. Tran, H.-N., Nguyen, Q.-H., Jeong, J., Loi, D.-L., Nam, Y.H., Kang, T.H., Yoon, J., Baek, K., and Jeong, Y. (2023). The embryonic patterning gene Dbx1 governs the survival of the auditory midbrain via Tcf7l2-Ap2δ transcriptional cascade. Cell Death Differ. 30, 1563–1574. 10.1038/s41418-023-01165-6.

41. Mouse Genome Informatics (MGI) Tg(Ntsr1-cre)GN209Gsat transgene detail (MGI:4367043). (The Jackson Laboratory). The Jackson Laboratory The Jackson Laboratory.

42. Gonzalez, D., Tomasek, M., Hays, S., Sridhar, V., Ammanuel, S., Chang, C., Pawlowski, K., Huber, K.M., and Gibson, J.R. (2019). Audiogenic Seizures in the *Fmr1* Knock-Out Mouse Are Induced by *Fmr1* Deletion in Subcortical, VGlut2-Expressing Excitatory Neurons and Require Deletion in the Inferior Colliculus. J. Neurosci. 39, 9852–9863. 10.1523/JNEUROSCI.0886-19.2019.

43. GENSAT. (2003). Cre images for neurotensin receptor 1 (Ntsr1), founder line GN209. http://www.gensat.org/creGeneView.jsp?founder_id=44880&gene_id=511&backcrossed=false http://www.gensat.org/creGeneView.jsp?founder_id=44880&gene_id=511&backcrossed=false.

44. Gómez-Nieto, R., Hormigo, S., and López, D.E. (2020). Prepulse Inhibition of the Auditory Startle Reflex Assessment as a Hallmark of Brainstem Sensorimotor Gating Mechanisms. Brain Sci. 10, 639. 10.3390/brainsci10090639.

45. Land, R., Burghard, A., and Kral, A. (2016). The contribution of inferior colliculus activity to the auditory brainstem response (ABR) in mice. Hear. Res. 341, 109–118. 10.1016/j.heares.2016.08.008.

46. Cramer, K.S., and Miko, I.J. (2016). Eph-ephrin signaling in nervous system development. F1000Research 5, 413. 10.12688/f1000research.7417.1.

47. Wallace, M.M., Harris, J.A., Brubaker, D.Q., Klotz, C.A., and Gabriele, M.L. (2016). Graded and discontinuous EphA–ephrinB expression patterns in the developing auditory brainstem. Hear. Res. 335, 64–75. 10.1016/j.heares.2016.02.013.

48. Talarico, M., De Bellescize, J., De Wachter, M., Le Guillou, X., Le Meur, G., Egloff, M., Isidor, B., Cogné, B., Beysen, D., Rollier, P., et al. (2025). RORA-neurodevelopmental disorder: A unique triad of developmental disabilities, cerebellar anomalies, and myoclonic seizures. Genet. Med. 27, 101347. 10.1016/j.gim.2024.101347.

49. Li, S., Zhou, D., Han, Y., Feng, C., Lv, X., Fu, G., Wei, X., Wu, C., Yu, C., and Yang, Z. (2025). Genetic and functional analysis of ZMIZ1 in neurodevelopmental disorder with dysmorphic facies and distal skeletal anomalies (NEDDFSA): insights from muscle cells and signaling pathways. Pediatr. Res. 10.1038/s41390-025-04612-x.

50. Tomioka, R., Kobayashi, K., and Song, W.-J. (2026). Anatomical and neurochemical profiles of somatostatin-positive neurons in the mouse inferior colliculus. Neuroscience, S0306452226001600. 10.1016/j.neuroscience.2026.03.004.

51. Liu, M., Hu, Q., Feng, W., Guo, Q., Ma, T., Li, X., Meng, Q., Xu, S., Li, J., Zhang, T., et al. (2026). Molecular taxonomy and spatial organization define neuronal subtypes in the mouse inferior colliculus. iScience 29, 116484. 10.1016/j.isci.2026.116484.

52. Spiteri, E., Konopka, G., Coppola, G., Bomar, J., Oldham, M., Ou, J., Vernes, S.C., Fisher, S.E., Ren, B., and Geschwind, D.H. (2007). Identification of the Transcriptional Targets of FOXP2, a Gene Linked to Speech and Language, in Developing Human Brain. Am. J. Hum. Genet. 81, 1144–1157. 10.1086/522237.

53. Griswold, A.J., Ma, D., Cukier, H.N., Nations, L.D., Schmidt, M.A., Chung, R.-H., Jaworski, J.M., Salyakina, D., Konidari, I., Whitehead, P.L., et al. (2012). Evaluation of copy number variations reveals novel candidate genes in autism spectrum disorder-associated pathways. Hum. Mol. Genet. 21, 3513–3523. 10.1093/hmg/dds164.

54. Koromina, M., Flitton, M., Blockley, A., Mellor, I.R., and Knight, H.M. (2019). Damaging coding variants within kainate receptor channel genes are enriched in individuals with schizophrenia, autism and intellectual disabilities. Sci. Rep. 9, 19215. 10.1038/s41598-019-55635-4.

55. Wamsley, B., Bicks, L., Cheng, Y., Kawaguchi, R., Quintero, D., Margolis, M., Grundman, J., Liu, J., Xiao, S., Hawken, N., et al. (2024). Molecular cascades and cell type–specific signatures in ASD revealed by single-cell genomics. Science 384, eadh2602. 10.1126/science.adh2602.

56. Mayfield, J., Blednov, Y.A., and Harris, R.A. (2015). Behavioral and Genetic Evidence for GIRK Channels in the CNS. In International Review of Neurobiology (Elsevier), pp. 279–313. 10.1016/bs.irn.2015.05.016.

57. Marron Fernandez De Velasco, E., Zhang, L., N. Vo, B., Tipps, M., Farris, S., Xia, Z., Anderson, A., Carlblom, N., Weaver, C.D., Dudek, S.M., et al. (2017). GIRK2 splice variants and neuronal G protein-gated K+ channels: implications for channel function and behavior. Sci. Rep. 7, 1639. 10.1038/s41598-017-01820-2.

58. Schmunk, G., and Gargus, J.J. (2013). Channelopathy pathogenesis in autism spectrum disorders. Front. Genet. 4. 10.3389/fgene.2013.00222.

59. Krüger, J., Schubert, J., Kegele, J., Labalme, A., Mao, M., Heighway, J., Seebohm, G., Yan, P., Koko, M., Aslan-Kara, K., et al. (2022). Loss-of-function variants in the KCNQ5 gene are implicated in genetic generalized epilepsies. eBioMedicine 84, 104244. 10.1016/j.ebiom.2022.104244.

60. Zhang, Y., Tachtsidis, G., Schob, C., Koko, M., Hedrich, U.B.S., Lerche, H., Lemke, J.R., Van Haeringen, A., Ruivenkamp, C., Prescott, T., et al. (2021). *KCND2* variants associated with global developmental delay differentially impair Kv4.2 channel gating. Hum. Mol. Genet. 30, 2300–2314. 10.1093/hmg/ddab192.

61. Nabit, B.P., Taylor, A., and Winder, D.G. (2024). Thalamocortical mGlu8 Modulates Dorsal Thalamus Excitatory Transmission and Sensorimotor Activity. J. Neurosci. 44, e0119242024. 10.1523/JNEUROSCI.0119-24.2024.

62. Shen, K., and Johnson, S.W. (2003). Group II metabotropic glutamate receptor modulation of excitatory transmission in rat subthalamic nucleus. J. Physiol. 553, 489–496. 10.1113/jphysiol.2003.052209.

63. Breiderhoff, T., Christiansen, G.B., Pallesen, L.T., Vaegter, C., Nykjaer, A., Holm, M.M., Glerup, S., and Willnow, T.E. (2013). Sortilin-Related Receptor SORCS3 Is a Postsynaptic Modulator of Synaptic Depression and Fear Extinction. PLoS ONE 8, e75006. 10.1371/journal.pone.0075006.

64. Savas, J.N., Ribeiro, L.F., Wierda, K.D., Wright, R., DeNardo-Wilke, L.A., Rice, H.C., Chamma, I., Wang, Y.-Z., Zemla, R., Lavallée-Adam, M., et al. (2015). The Sorting Receptor SorCS1 Regulates Trafficking of Neurexin and AMPA Receptors. Neuron 87, 764–780. 10.1016/j.neuron.2015.08.007.

65. The DDD Study, Homozygosity Mapping Collaborative for Autism, UK10K Consortium, The Autism Sequencing Consortium, De Rubeis, S., He, X., Goldberg, A.P., Poultney, C.S., Samocha, K., Ercument Cicek, A., et al. (2014). Synaptic, transcriptional and chromatin genes disrupted in autism. Nature 515, 209–215. 10.1038/nature13772.

66. Li, L., Korngut, L.M., Frost, B.J., and Beninger, R.J. (1998). Prepulse inhibition following lesions of the inferior colliculus: prepulse intensity functions. Physiol. Behav. 65, 133–139. 10.1016/S0031-9384(98)00143-7.

67. Kurt, S., Groszer, M., Fisher, S.E., and Ehret, G. (2009). Modified sound-evoked brainstem potentials in Foxp2 mutant mice. Brain Res. 1289, 30–36. 10.1016/j.brainres.2009.06.092.

68. Buck, A., Dupont, T., Andrews Cavanagh, R., Postal, O., Bourien, J., Puel, J., Michalski, N., and Gourévitch, B. (2025). Neural Response Reliability as a Marker of the Transition of Neural Codes along Auditory Pathways. Adv. Sci. 12, e08777. 10.1002/advs.202508777.

69. Iwata, R., Casimir, P., Erkol, E., Boubakar, L., Planque, M., Gallego López, I.M., Ditkowska, M., Gaspariunaite, V., Beckers, S., Remans, D., et al. (2023). Mitochondria metabolism sets the species-specific tempo of neuronal development. Science 379, eabn4705. 10.1126/science.abn4705.

70. Zeller, K., Rahner-Welsch, S., and Kuschinsky, W. (1997). Distribution of Glut1 Glucose Transporters in Different Brain Structures Compared to Glucose Utilization and Capillary Density of Adult Rat Brains. J. Cereb. Blood Flow Metab. 17, 204–209. 10.1097/00004647-199702000-00010.

71. Sarachana, T., and Hu, V.W. (2013). Genome-wide identification of transcriptional targets of RORA reveals direct regulation of multiple genes associated with autism spectrum disorder. Mol. Autism 4, 14. 10.1186/2040-2392-4-14.

72. BenÃ-tez-Burraco, A., and Boeckx, C. (2014). FOXP2, retinoic acid, and language: a promising direction. Front. Cell. Neurosci. 8. 10.3389/fncel.2014.00387.

73. Konopka, G., Bomar, J.M., Winden, K., Coppola, G., Jonsson, Z.O., Gao, F., Peng, S., Preuss, T.M., Wohlschlegel, J.A., and Geschwind, D.H. (2009). Human-specific transcriptional regulation of CNS development genes by FOXP2. Nature 462, 213–217. 10.1038/nature08549.

74. Karayannis, T., Au, E., Patel, J.C., Kruglikov, I., Markx, S., Delorme, R., Héron, D., Salomon, D., Glessner, J., Restituito, S., et al. (2014). Cntnap4 differentially contributes to GABAergic and dopaminergic synaptic transmission. Nature 511, 236–240. 10.1038/nature13248.

75. Wang, C., Pan, Y.-H., Wang, Y., Blatt, G., and Yuan, X.-B. (2019). Segregated expressions of autism risk genes Cdh11 and Cdh9 in autism-relevant regions of developing cerebellum. Mol. Brain 12, 40. 10.1186/s13041-019-0461-4.

76. Shimoyama, Y., Tsujimoto, G., Kitajima, M., and Natori, M. (2000). Identification of three human type-II classic cadherins and frequent heterophilic interactions between different subclasses of type-II classic cadherins.

77. DDD study, Verheije, R., Kupchik, G.S., Isidor, B., Kroes, H.Y., Lynch, S.A., Hawkes, L., Hempel, M., Gelb, B.D., Ghoumid, J., et al. (2019). Heterozygous loss-of-function variants of MEIS2 cause a triad of palatal defects, congenital heart defects, and intellectual disability. Eur. J. Hum. Genet. 27, 278–290. 10.1038/s41431-018-0281-5.

78. Hizawa, K., Sasaki, T., and Arimura, N. (2024). A comparative overview of DSCAM and its multifunctional roles in Drosophila and vertebrates. Neurosci. Res. 202, 1–7. 10.1016/j.neures.2023.12.005.

79. Gene Expression Nervous System Atlas (GENSAT) GENSAT. Cre images for neurotensin receptor 1 (Ntsr1), founder line GN209. (Rockefeller University). http://www.gensat.org/creGeneView.jsp?founder_id=44880&gene_id=511&backcrossed=false&utm_source=chatgpt.com http://www.gensat.org/creGeneView.jsp?founder_id=44880&gene_id=511&backcrossed=false&utm_source=chatgpt.com.

80. Gong, S., Doughty, M., Harbaugh, C.R., Cummins, A., Hatten, M.E., Heintz, N., and Gerfen, C.R. (2007). Targeting Cre Recombinase to Specific Neuron Populations with Bacterial Artificial Chromosome Constructs. J. Neurosci. 27, 9817–9823. 10.1523/JNEUROSCI.2707-07.2007.

81. Lovelace, J.W., Wen, T.H., Reinhard, S., Hsu, M.S., Sidhu, H., Ethell, I.M., Binder, D.K., and Razak, K.A. (2016). Matrix metalloproteinase-9 deletion rescues auditory evoked potential habituation deficit in a mouse model of Fragile X Syndrome. Neurobiol. Dis. 89, 126–135. 10.1016/j.nbd.2016.02.002.

82. Lovelace, J.W., Ethell, I.M., Binder, D.K., and Razak, K.A. (2018). Translation-relevant EEG phenotypes in a mouse model of Fragile X Syndrome. Neurobiol. Dis. 115, 39–48. 10.1016/j.nbd.2018.03.012.

83. Niell, C.M., and Stryker, M.P. (2010). Modulation of Visual Responses by Behavioral State in Mouse Visual Cortex. Neuron 65, 472–479. 10.1016/j.neuron.2010.01.033.

84. Holley, A.J., Shedd, A., Boggs, A., Lovelace, J., Erickson, C., Gross, C., Jankovic, M., Razak, K., Huber, K., and Gibson, J.R. (2022). A sound-driven cortical phase-locking change in the Fmr1 KO mouse requires Fmr1 deletion in a subpopulation of brainstem neurons. Neurobiol. Dis. 170, 105767. 10.1016/j.nbd.2022.105767.

85. Maris, E., and Oostenveld, R. (2007). Nonparametric statistical testing of EEG- and MEG-data. J. Neurosci. Methods 164, 177–190. 10.1016/j.jneumeth.2007.03.024.

86. Lovelace, J.W., Rais, M., Palacios, A.R., Shuai, X.S., Bishay, S., Popa, O., Pirbhoy, P.S., Binder, D.K., Nelson, D.L., Ethell, I.M., et al. (2020). Deletion of Fmr1 from Forebrain Excitatory Neurons Triggers Abnormal Cellular, EEG, and Behavioral Phenotypes in the Auditory Cortex of a Mouse Model of Fragile X Syndrome. Cereb. Cortex 30, 969–988. 10.1093/cercor/bhz141.

87. Khandelwal, N., Cavalier, S., Rybalchenko, V., Kulkarni, A., Anderson, A.G., Konopka, G., and Gibson, J.R. (2021). FOXP1 negatively regulates intrinsic excitability in D2 striatal projection neurons by promoting inwardly rectifying and leak potassium currents. Mol. Psychiatry 26, 1761–1774. 10.1038/s41380-020-00995-x.

88. Ortiz, A., Ayhan, F., Khandelwal, N., Outland, E., Jankovic, M., Harper, M., and Konopka, G. (2025). Cell-type-specific roles of FOXP1 in the excitatory neuronal lineage during early neocortical murine development. Cell Rep. 44, 115384. 10.1016/j.celrep.2025.115384.

89. Yao, Z., Van Velthoven, C.T.J., Kunst, M., Zhang, M., McMillen, D., Lee, C., Jung, W., Goldy, J., Abdelhak, A., Aitken, M., et al. (2023). A high-resolution transcriptomic and spatial atlas of cell types in the whole mouse brain. Nature 624, 317–332. 10.1038/s41586-023-06812-z.

90. MapMyCells (Allen Institute for Brain Science). https://portal.brain-map.org/atlases-and-data/bkp/mapmycells

